# Fast and Accurate Bayesian Polygenic Risk Modeling with Variational Inference

**DOI:** 10.1101/2022.05.10.491396

**Authors:** Shadi Zabad, Simon Gravel, Yue Li

**Affiliations:** School of Computer Science, McGill University; Department of Human Genetics, McGill University

## Abstract

The recent proliferation of large scale genome-wide association studies (GWASs) has motivated the development of statistical methods for phenotype prediction using single nucleotide polymorphism (SNP) array data. These polygenic risk score (PRS) methods formulate the task of polygenic prediction in terms of a multiple linear regression framework, where the goal is to infer the joint effect sizes of all genetic variants on the trait. Among the subset of PRS methods that operate on GWAS summary statistics, sparse Bayesian methods have shown competitive predictive ability. However, most existing Bayesian approaches employ Markov Chain Monte Carlo (MCMC) algorithms for posterior inference, which are computationally inefficient and do not scale favorably with the number of SNPs included in the analysis. Here, we introduce Variational Inference of Polygenic Risk Scores (`VIPRS`), a Bayesian summary statistics-based PRS method that utilizes Variational Inference (VI) techniques to efficiently approximate the posterior distribution for the effect sizes. Our experiments with genome-wide simulations and real phenotypes from the UK Biobank (UKB) dataset demonstrated that variational approximations to the posterior are competitively accurate and highly efficient. When compared to state-of-the-art PRS methods, `VIPRS` consistently achieves the best or second best predictive accuracy in our analyses of 36 simulation configurations as well as 12 real phenotypes measured among the UKB participants of “White British” background. This performance advantage was higher among individuals from other ethnic groups, with an increase in *R*^2^ of up to 1.7-fold among participants of Nigerian ancestry for Low-Density Lipoprotein (LDL) cholesterol. Furthermore, given its computational efficiency, we applied `VIPRS` to a dataset of up to 10 million genetic markers, an order of magnitude greater than the standard HapMap3 subset used to train existing PRS methods. Modeling this expanded set of variants conferred significant improvements in prediction accuracy for a number of highly polygenic traits, such as standing height.

## 1 Introduction

In recent years, with the rapid growth of large-scale biobank data with comprehensive genotyping and phenotyping efforts [1–3], there has been growing interest in developing statistical methods to quantify an individual’s disease risk from their genotype data [4–8]. At the same time, these rich biobank data sources have powered many recent analyses of complex traits and diseases, revealing highly polygenic architectures [9–11] with a wide range of effect sizes across different genomic categories [12–14]. Linear models are an important framework for complex trait analysis which allow for the estimation of the additive genetic component of a phenotype, also known as a polygenic score (PGS) or polygenic risk score (PRS) in clinical contexts [5, 15]. Even though many examples of genetic interactions have been documented, such additive effects capture much of the genetic variation underlying human complex traits [16, 17]. Recent work has highlighted the clinical relevance of polygenic scores for some diseases and health conditions [18, 19], especially in applications related to disease risk stratification [20–22] and personalized medicine [23].

Estimating polygenic scores from genome-wide association study (GWAS) data has been an active area of research, with numerous methods recently developed [4, 6, 24–32]. Standard PRS methods formulate the problem of polygenic risk estimation in terms of a multiple linear regression framework, where the goal is to infer the joint effect sizes of all genetic variants on the trait. The most common class of genetic variation considered in these analyses are single nucleotide polymorphisms (SNPs), which are either measured by modern genotyping arrays or statistically imputed using reference haplotypes [33, 34].

Genotyping arrays combined with imputation can accurately capture the genotype of an individual at millions of genetic markers. When paired with modern GWAS sample sizes routinely exceeding hundreds of thousands of individuals, high dimensional data of this scale present several computational and statistical challenges. Furthermore, most individual-level GWAS data sources are protected for privacy concerns [35]. These two factors motivated the development of a number of PRS methods that estimate polygenic risk scores based on GWAS summary statistics alone [4, 6, 25–27, 29–32], which are the marginal test statistic per SNP.

Within this class of summary statistics-based methods, Bayesian PRS models enable a principled way to incorporate prior knowledge as probability distributions over the genetic causal architecture of complex traits. In addition to providing meaningful estimates of parameter uncertainties [36], Bayesian approaches have shown competitive predictive ability, exceeding the predictive performance of heuristic or penalized estimators in many settings [6, 26, 30, 31, 37, 38]. However, a major limitation of some existing Bayesian methods is that their scalability is hampered by slow and inefficient inference techniques. While heuristic methods such as Clumping-and-Thresholding (C+T) are routinely applied on millions of SNPs, Bayesian approaches are generally restricted to a smaller subset of approximately one million genetic markers. One of the main reasons for this limitation stems from computational considerations: Most Bayesian PRS methods employ Markov Chain Monte Carlo (MCMC) algorithms to approximate the posterior for the effect sizes [4, 6, 26, 30]. MCMC algorithms are known to be asymptotically accurate but often slow to converge [39, 40]. In practice, to obtain accurate posterior estimates, the MCMC chains need to be run for hundreds or thousands of iterations [4, 6]. This challenge can be partially remedied with the help of efficient software implementation and enhanced linear algebra routines, which recently enabled scaling up two well-known MCMC-based Bayesian PRS methods to several million SNPs [6, 38]. While this is an important advance, these variants still constitute a small fraction of the genetic variation that can be assayed by modern whole genome-sequencing technologies [3, 41].

An alternative scheme for approximating the posterior density for the effect sizes is Variational Inference (VI), a fast and deterministic class of algorithms that recast the problem of posterior inference in the form of an optimization problem [39, 40, 42, 43]. Variational methods have seen a surge of interest in the machine learning literature in recent years due to significant advances in stochastic optimization techniques [44, 45]. Methods that utilize variational inference have been explored in a wide variety of statistical genetics applications, specifically in the context of linear mixed models (LMMs) [46], association mapping [47, 48], fine-mapping [49, 50], and enrichment analysis [51, 52], among others. More recently, a number of studies examined the properties and relative accuracy of certain variational approximations to PRS using both individual level data and summary statistics [32, 53–55].

In this work, we present `VIPRS`, a Bayesian summary statistics-based PRS method that utilizes variational inference to approximate the posterior for the effect sizes. We conduct a comprehensive set of experiments using simulated and real traits to assess the predictive ability of `VIPRS` in comparison with the some of the most popular Bayesian and non-Bayesian PRS methods. Overall, we show that `VIPRS` is a scalable and flexible method that enjoys the speed and efficiency of heuristic approaches such as Clumping-and-Thresholding (C+T), while rivaling state of the art Bayesian methods in terms of its predictive performance. We demonstrate the flexibility of the method by testing its predictions with different families of priors on the effect size, paired with four distinct strategies for tuning the hyperparameters of the model. To illustrate its scalability, we evaluate the predictive accuracy of `VIPRS` with approximately 10 million SNPs, almost an order of magnitude greater than the standard HapMap3 subset routinely used for this task. This enables us to examine the potential for phenotype prediction of a number highly polygenic traits at a much finer scale.

## 2 Results

### 2.1 Variational inference of polygenic risk scores (VIPRS)

Given a random sample of individuals from a general population with paired genotype and phenotype data, represented by a genotype matrix **X**_*N*×*M*_ and a corresponding phenotype vector **y**_*N*×1_, we model the dependence of the phenotype on the genotype via the standard linear model: **y** = **Xβ** + ***ϵ***. Here, **β**_*M*×1_ is a random and unknown vector of effect sizes for each of the *M* variants included in the model and ***ϵ***_*N*×1_ is a vector of residuals. In this setup, the Bayesian formulation of polygenic risk modeling is concerned with inferring the posterior distribution for the effect sizes of the genetic variants on the trait,

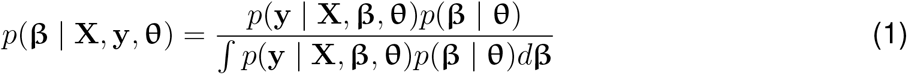

where **θ** is a composite term for the hyperparameters of the model, such as the residual variance 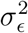. The posterior distribution for the effect sizes is proportional to the likelihood *p*(**y** | **X**, **β**, **θ**) and the prior *p*(**β** | **θ**), with the latter encapsulating prior beliefs about the genetic architecture of the trait. For continuous traits, assuming that the samples are unrelated and ancestrally homogeneous, we use the common model of the likelihood as an isotropic Gaussian distribution, 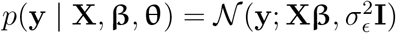.

As for the prior, a common assumption that has proved powerful in predictive settings [4, 30] is to model the effect size of each variant with a two-component sparse mixture distribution: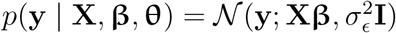. Under this prior distribution, with probability *π* the effect size is drawn from a Gaussian centered at zero with the scale proportional to *σ_β_*, and with probability 1 − *π* from a Dirac delta density centered at zero. The first component captures the contribution of causal variants whereas the second component models the role of variants with null effects. This is sometimes known as the spike-and-slab prior and is used to enforce sparsity on the effect size estimates [56–58]. In addition to the standard spike-and-slab prior density, we also explore implementations of the model with a sparse Gaussian mixture prior [6, 24, 55] as well as a frequency dependent prior, i.e. the Alpha prior (**Supplementary Information**). A major difficulty in this context is that these flexible families of priors result in an analytically intractable posterior density, due to the integral in the model evidence ∫ *p*(**y** | **X**, **β**, **θ**)*p*(**β** | **θ**)*d***β**. This requires the use of approximate posterior inference schemes.

Most existing methods for Bayesian PRS estimation are based on Markov Chain Monte Carlo (MCMC) inference algorithms, such as Gibbs sampling [4, 6, 26, 30]. Despite their many successful applications, MCMC methods can be computationally inefficient for performing inference over high-dimensional genome-wide data containing millions of SNPs. Additionally, with MCMC algorithms, it is difficult to assess model convergence [39, 40]. As an alternative, in this study, we present a method for Variational Inference of PRS (`VIPRS`). We approximate the posterior distribution of the effect sizes by a proposed parametric density, *q*(**β**, **s**), and optimize its parameters to minimize a global measure of the divergence from the true posterior [42, 43, 48, 49]. For efficient inference, we use a paired mean-field distribution family [49, 59] that factorizes the joint density of the effect sizes into the product of the individual densities for the effect size of each variant,

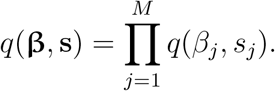

Here, *s_j_* is the posterior Bernoulli variable indicating whether variant *j* is causal for the trait of interest. Under this posterior variational family, the SNP effect sizes are assumed to be independent, a restrictive assumption that nonetheless works well in practice. In the **Methods** section and accompanying **Supplementary Information**, we show that this formulation results in a fast coordinate-ascent optimization procedure that, under some assumptions, can be expressed solely in terms of GWAS summary statistics. Related variational approaches have been explored in the context of fine mapping [49, 50] and polygenic modeling [32, 53, 55], but differ in both conceptual and technical details – we discuss these similarities and differences in **Supplementary Information**.

Inference under this model, however, requires setting the hyperparameters 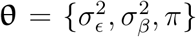, for which we explored four different strategies. In the basic formulation of the `VIPRS` model, we use a Variational Expectation-Maximization (VEM) framework, also known as empirical Bayes. In the E-Step, we update the parameters of the variational density *q*(**β**, **s**); in the M-Step, we update the hyperparameters using their maximum likelihood estimates [49, 60]. Additionally, we experimented with grid search (GS) (`VIPRS-GS`) [61], Bayesian optimization (BO) (`VIPRS-BO`) [62], and Bayesian Model Averaging (BMA) (`VIPRS-BMA`) [49] as means to set or integrate out some or all the hyperparameters of the model (**Methods**).

### 2.2 Genome-wide simulation results

To examine the predictive performance of the `VIPRS` model compared to existing PRS methods, we simulated quantitative and case-control traits with varying genetic architectures and heritability values. To align our simulations with the real trait analyses in terms of cohort size and composition, we used genotype data for a subset of ≈ 340, 000 unrelated White British individuals from the UK Biobank (**Methods**) and a HapMap3 subset of ≈ 1.1 million genotyped and imputed SNPs. The simulations followed the generative models outlined in the **Methods** section and **Supplementary Information**, with the effect size of each variant drawn from different architectures and residuals for each individual sampled from an isotropic Gaussian density. For binary traits, we simulated case-control status following the Liability Threshold model [63], with the prevalence set to 15%.

The simulations spanned six different genetic architectures, including both sparse and infinitesimal scenarios. In the first three scenarios, we simulated under the spike-and-slab model (**Methods**), where the effect size for a given variant *j* was drawn from 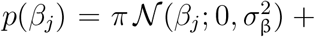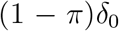, with the proportion of SNPs contributing to the variation in the trait ranging along a pre-specified grid *π* ∈ {10^−4^, 10^−3^, 10^−2^}. In the next two simulation scenarios, the effect size is drawn from a scale mixture of Gaussians, 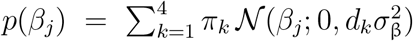, with the mixing proportions set to *π* = {0.95, 0.02, 0.02, 0.01}. The variance multipliers *d_k_* were set to *d* = {0.0, 0.01, 0.1, 1} for the sparse mixture model and *d* = {0.001, 0.01, 0.1, 1} for the infinitesimal mixture model. Finally, for the infinitesimal model, we assumed that the effect size for all variants is drawn from a zero-centered Gaussian density, 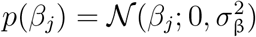. For each genetic architecture, we varied the proportion of additive genetic variance captured by all causal SNPs, 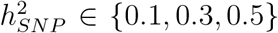, such that the simulated traits range from the mildly to the highly heritable. For each unique configuration, we simulated 10 independent phenotypes, for a total of 180 traits for each class (binary and continuous). Once the traits were simulated for all individuals in the dataset, we randomly split the sample into 70% training, 15% validation, and 15% testing, with the training set used to generate GWAS summary statistics.

Next, we fit the `VIPRS` model to the summary statistics from the training data, along with other commonly-used PRS methods. Given the many existing PRS approaches, we selected for comparison five methods that performed favorably in recent comprehensive surveys [37, 38, 64], namely `SBayesR` [6], `LDPred2` [30], `PRScs` [26], `MegaPRS` [32], and `Lassosum` [25]. The first three methods use the Bayesian framework outlined above for approximate posterior inference, all employing a Gibbs sampling algorithm. They are mainly distinguished by the families of prior density they assign to the effect sizes, among many other algorithmic choices. The fourth model, `MegaPRS`, is similar to `VIPRS` in that it uses variational inference for polygenic risk estimation, though there are some fundamental differences in the details of the optimization algorithm (see **Supplementary Information**). Finally, we included `PRSice2` [27], which implements clumping and thresholding, a commonly used baseline model. After fitting each method on the summary statistics from the training data, we used the effect size estimates to generate polygenic scores for individuals in the held-out test set and evaluated their predictive performance. For quantitative traits, we computed the incremental Prediction *R*^2^ for each model, while for binary traits we show the area under the precision-recall curve (AUPRC), a preferable metric in the presence of class imbalance [65]. In addition to the six external PRS models, we also examined the predictive performance of the basic `VIPRS` model trained with the Variational EM framework (**Methods**) as well as a version of the `VIPRS` model, dubbed `VIPRS-GS`, in which we perform grid search and tune the hyperparameters based on predictive performance on a held-out validation set.

The predictive performance results for this simulation study are summarized in **Fig.** 1 and **Supplementary Fig.** S1, which show that `VIPRS-GS` outperforms or is on-par with state-of-the-art PRS methods in most of the scenarios tested. In particular, our analyses indicate that `VIPRS` provides the most benefit for more sparse architectures and highly heritable traits (left-most panels in **Fig.** 1(a) and **Supplementary Fig.** S1(a)). Notably, in this particular setting, `VIPRS` is able to capture most of the additive genetic variance (as measured by the *R*^2^ metric, which is upper-bounded by the heritability), while other Bayesian and non-Bayesian methods often lag behind. For infinitesimal and mixture-based architectures, `VIPRS` shows competitive predictive ability across the range, only lagging slightly behind `SBayesR` in those settings. For highly polygenic traits with the proportion of causal variants equal or greater than 1%, all models conferred lower prediction accuracy relative the heritability values that we simulated with. This is because, under our simplified simulation scenario, the larger the number of causal variants, the smaller the effect size per SNP. Consequently, this makes it more difficult for PRS methods to pick up the true causal signals, at least given the training sample sizes available. Nonetheless, the `VIPRS` models conferred higher predictive performance relative to most competing methods in many of those scenarios. This pattern holds for both quantitative (**Fig.** 1) as well as binary case-control phenotypes (**Supplementary Fig.** S1). This improvement in prediction accuracy comes also with improved computational efficiency, with the run-time of the standard `VIPRS` model rivaling other heuristic and deterministic methods, such as `Lassosum`, `MegaPRS`, and `PRSice2` (**Fig.** 3).

**Figure 1:**
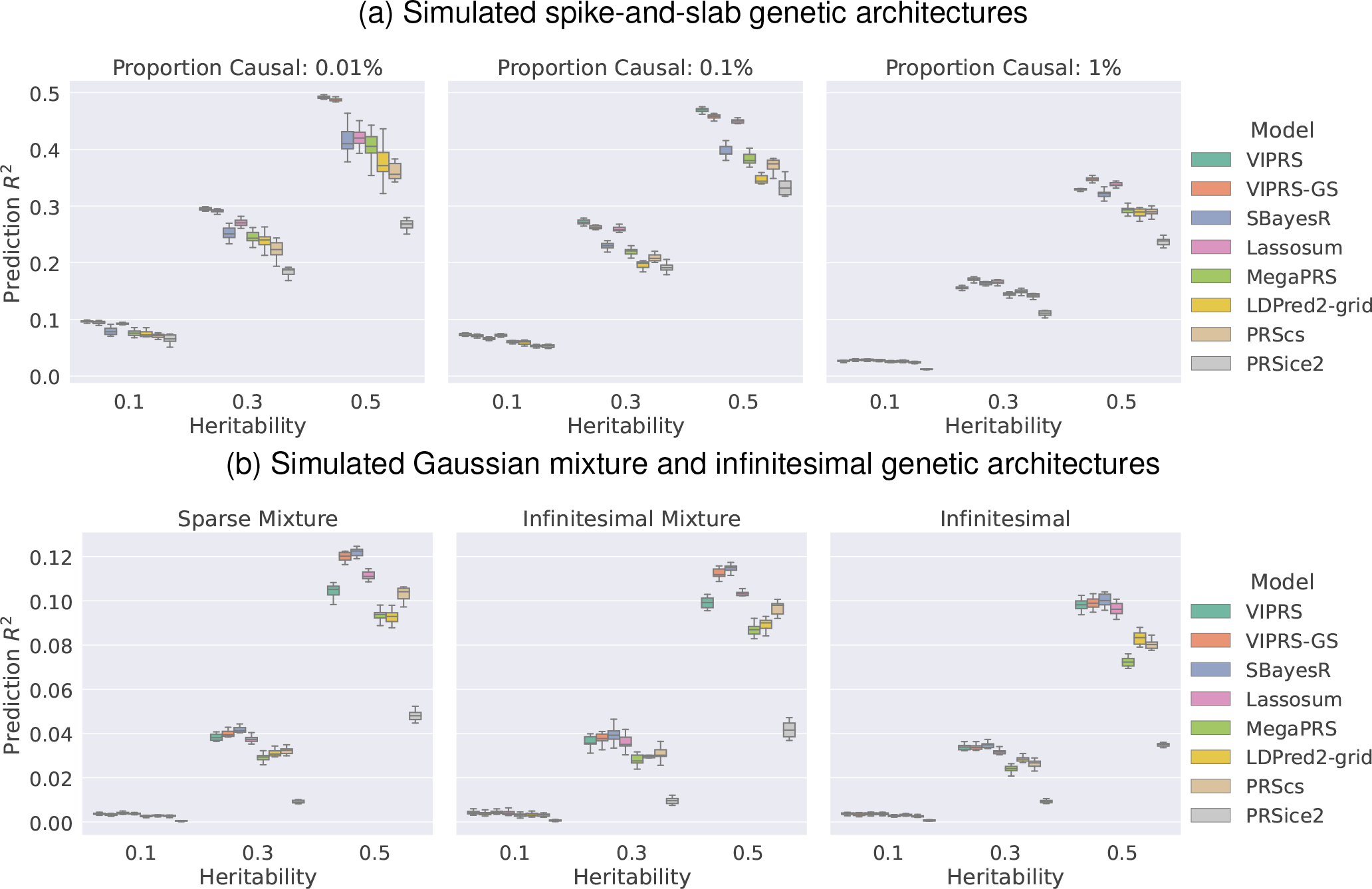
Predictive performance of summary statistics-based PRS methods on simulated quantitative traits following **(a)** spike-and-slab and **(b)** Gaussian mixture or infinitesimal genetic architectures. The phenotypes were simulated using real genotype data from the White British cohort in the UK Biobank (*N* = 337, 225), leveraging a subset of 1.1 million HapMap3 variants. The simulation scenarios encompass a total of 18 configurations, spanning 6 genetic architectures and 3 values for SNP heritability. For each configuration, we simulated 10 independent phenotypes. Each panel shows results for phenotypes simulated with the pre-specified genetic architecture and each column within a panel shows performance metrics for phenotypes simulated with a pre-specified SNP heritability. The performance metric shown is the incremental prediction *R*^2^. The boxplot for each method and simulation configuration shows the quartiles of the *R*^2^ scores for the 10 simulated phenotypes. The PRS methods shown are our proposed `VIPRS` and `VIPRS-GS` (using grid search to tune model hyperparameters) as well as 5 other baseline models: `SBayesR`, `Lassosum`, `MegaPRS`, `LDPred2` (grid), `PRScs`, and `PRSice2` (C+T).

The prediction accuracy of PRS methods on simulated phenotypes may be over-optimistic due the similarity between the generative process for the simulations and their model assumptions. Therefore, it is important to systematically evaluate these methods on real phenotypes as shown next.

### 2.3 Application on real phenotypes in the UK Biobank

Given its competitive performance on simulated traits, we next sought to assess the relative predictive ability of the `VIPRS` model on real phenotypes measured for a subset of ≈ 340, 000 unrelated White British individuals in the UK Biobank. This focus on a large sub-cohort of relatively uniform ancestry helps us achieve sufficient power while reducing confounding due to population structure. A downside of this approach is that it is expected to yield PRS estimates that perform more poorly for individuals of other ancestries [66, 67], a limitation that we examine in more detail in the next section.

For this analysis, we extracted and processed phenotypic measurements for 9 quantitative traits and 3 binary traits that are commonly used to benchmark PRS methods (**Table** 1). The traits considered have varying (inferred) genetic architectures and SNP heritabilities. To make full use of the data, we followed a 5-fold cross-validation study design, where in each iteration, 80% of the samples with trait measurements were used to generate the GWAS summary statistics and training the PRS models and 20% were used as an independent test set.

**Table 1:** The list of real phenotypes and GWAS data sources analyzed in this study. With each phenotype code, we provide the full name and description, the GWAS data source (UKB or external), as well as the sample sizes for the training, validation, and test sets. The sample sizes for each subset may vary slightly across the 5 folds. We prepended some of the phenotype codes with `PASS` to indicate that the GWAS summary statistics are external to the UK Biobank and are publicly available. For analyses with the external GWAS summary statistics, the validation and test sets come from the UK Biobank.

| Phenotype | Description | GWAS Source | GWAS sample size | Validation sample size | Test sample size |
| --- | --- | --- | --- | --- | --- |
| HEIGHT | Standing height | UKB | 242,213 | 26,913 | 67,282 |
| HDL | high-density lipoprotein | UKB | 211,856 | 23,540 | 58,849 |
| BMI | Body mass index | UKB | 241,959 | 26,885 | 67,211 |
| FVC | Forced Vital Capacity | UKB | 221,249 | 24,584 | 61,459 |
| FEV1 | Forced Expiratory Volume in 1 second | UKB | 221,265 | 24,586 | 61,463 |
| HC | Hip circumference | UKB | 242,311 | 26,924 | 67,309 |
| WC | Waist circumference | UKB | 242,340 | 26,927 | 67,317 |
| LDL | low-density lipoprotein | UKB | 230,995 | 25,667 | 64,166 |
| BW | Birth weight | UKB | 138,300 | 15,367 | 38,417 |
| ASTHMA | Asthma | UKB | 229,031 | 25,448 | 63,620 |
| T2D | Type 2 diabetes | UKB | 235,937 | 26,216 | 65,538 |
| RA | Rheumatoid arthritis | UKB | 186,239 | 20,694 | 51,734 |
| PASS_HEIGHT | Standing height | [78] | 131,547 | 26,913 | 67,282 |
| PASS_HDL | High-density lipoprotein | [79] | 97,749 | 23,540 | 58,849 |
| PASS_BMI | Body mass index | [80] | 122,033 | 26,885 | 67,211 |
| PASS_LDL | Low-density lipoprotein | [79] | 93,354 | 25,667 | 64,166 |
| PASS_T2D | Type 2 diabetes | [81] | 60,786 | 26,216 | 65,538 |
| PASS_RA | Rheumatoid arthritis | [82] | 37,681 | 20,694 | 51,734 |

Our experiments show that, across a variety of different phenotypes, `VIPRS` is competitive with commonly-used Bayesian PRS methods (**Fig.** 2). Within the category of Bayesian PRS methods, the predictive performance of `VIPRS` is especially distinguished for anthropometric and blood lipid traits (**Fig.** 2(a)). For instance, when compared to the `LDPred2` model, which imposes the same spike-and-slab prior on the effect sizes, `VIPRS` shows an average of 4.6% improvement in prediction *R*^2^ on continuous traits. However, in many cases the basic `VIPRS` model lags behind the `SBayesR` [6] and `Lassosum` [25] models (**Fig.** 2). In addition to the difference in posterior approximation strategy (variational inference versus Gibbs sampling), the `SBayesR` model differs from the `VIPRS` model in three other important respects: (1) The prior on the effect size, (2) The estimator for the Linkage Disequilibrium (LD) between variants, and finally (3) the approach for estimating the hyperparameters of the model. We sought to understand the effect of each of these modeling choices on the predictive performance of our model.

**Figure 2:**
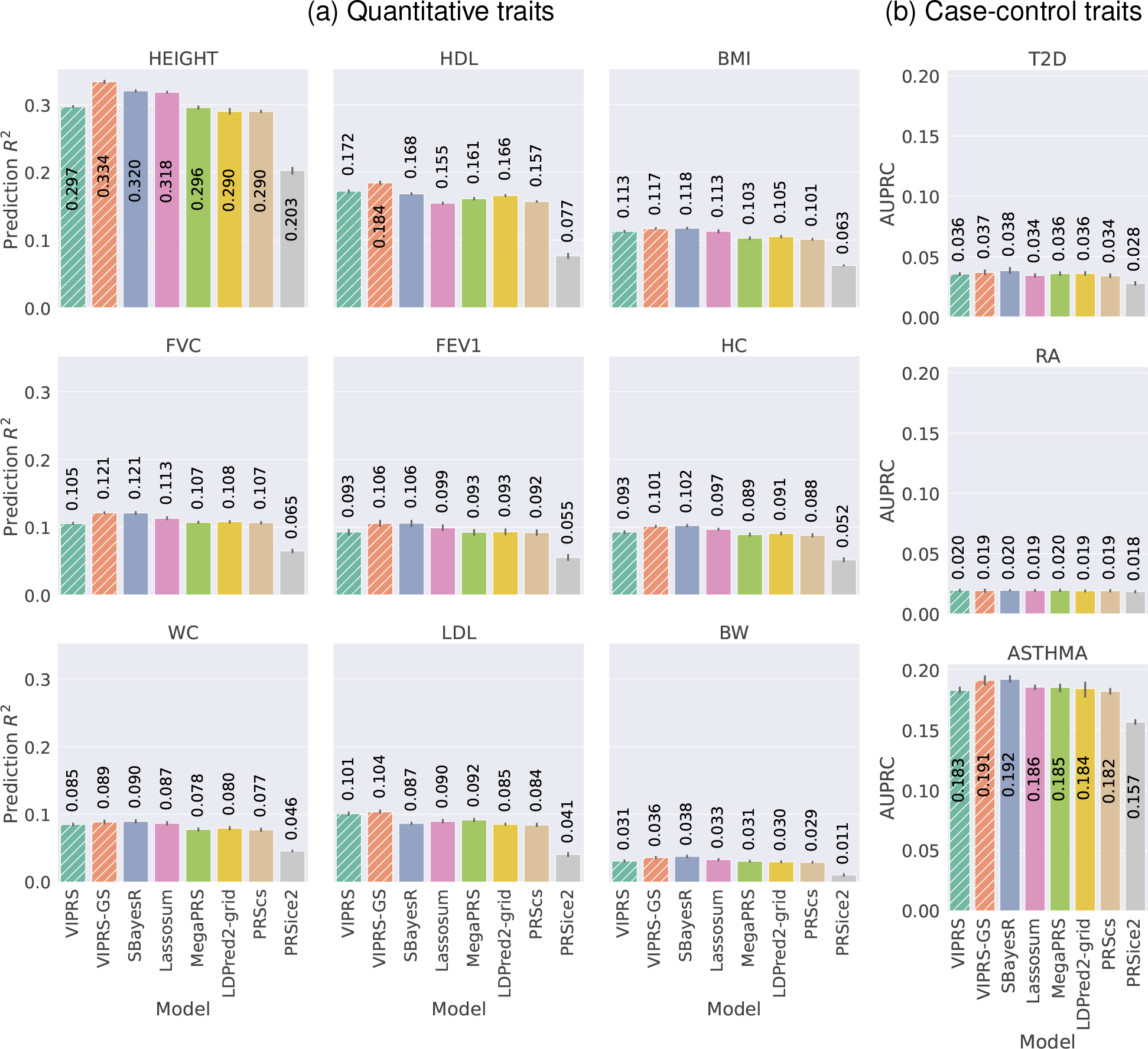
Predictive performance of summary statistics-based PRS methods on real **(a)** quantitative and **(b)** case-control phenotypes in the UK Biobank. The measured phenotypes were pre-processed and analyzed in a 5-fold cross-validation study design and the prediction metrics show the performance of each PRS method in predicting the phenotype in a held-out test set. Each panel shows the predictive performance, in terms of **(a)** incremental *R*^2^ and **(b)** area under the Precision Recall curve (AUPRC), of various PRS methods when applied to a given phenotype. The bars show the mean of the prediction metrics across the 5 folds and the black lines show the corresponding standard errors. The quantitative phenotypes analyzed are standing height (HEIGHT), high-density lipoprotein (HDL), body mass index (BMI), forced vital capacity (FVC), forced expiratory volume in 1 second (FEV1), hip circumference (HC), waist circumference (WC), low-density lipoprotein (LDL) and birth weight (BW). The binary phenotypes analyzed are asthma (ASTHMA), type 2 diabetes (T2D) and Rheumatoid arthritis (RA). The PRS methods shown are our proposed `VIPRS` and `VIPRS-GS` (using grid search to tune model hyperparameters) as well as 5 other baseline models: `SBayesR`, `Lassosum`, `MegaPRS`, `LDPred2` (grid), `PRScs`, and `PRSice2` (C+T). Dashed lines highlight the models contributed in this work.

To address the first point, we derived and implemented a version of `VIPRS` called `VIPRSMix`, where we replaced the spike-and-slab prior on the effect sizes with a sparse Gaussian mixture prior with four mixture components (**Supplementary Information**) [6, 24, 48]. Our experiments show that the more expressive mixture prior improves the performance of the standard `VIPRS` model on some traits, especially highly heritable and polygenic traits such as standing height and HDL (**Supplementary Fig.** S5), with on average of 2.4% increase in prediction *R*^2^ on continuous traits. However, the improvement is not consistent across all traits and using this prior does not fully bridge the gap between `VIPRS` and `SBayesR`.

Secondly, we assessed the impact of the LD estimator on the predictive performance of the `VIPRS` model by re-fitting the model with three commonly-used estimators for LD: windowed [30, 68], shrinkage [6, 69], and block [26, 70] (**Methods**). Our experiments indicate that, on many of the traits tested, using the shrinkage estimator for LD results in slight improvements in prediction accuracy, though, as in the case of the windowed LD estimator, it still slightly lags behind the `SBayesR` model (**Supplementary Fig.** S2, S3). Notably, however, the shrinkage estimator tends to be more robust when the sample size of the LD reference panel is small (**Supplementary Fig.** S2, S3).

Finally, and most importantly, the standard `VIPRS` model differs from the `SBayesR` model in terms of its hyperparameter estimation strategy. `SBayesR` follows a fully Bayesian approach for learning the hyperparameters of the model, assigning them priors and inferring their posterior distributions [6]. By contrast, `VIPRS` follows a Variational EM algorithm where in the M-Step we set the hyperparameters to their maximum-likelihood estimates [48, 49]. This latter strategy is known to be prone to overfitting or entrapment in local maxima [48, 49, 60, 71, 72]. As an alternative to the VEM framework, we tested three other strategies for tuning the hyperparameters of the model, including grid search [61], Bayesian optimization [62], and Bayesian model averaging [49] (see **Methods**, **Supplementary Fig.** S6,S7). In this context, similar to the `Lassosum`, `MegaPRS`, and `LDPred2` methods, we found that by setting some of the hyperparameters of the model via grid search with an independent validation set, `VIPRS-GS` provides a powerful remedy in most settings (**Fig.** 2, **Supplementary Fig.** S6,S7), resulting in a balanced trade-off between computational speed and predictive accuracy (**Fig.** 2 and 3). Indeed, our results show that the `VIPRS-GS` model conferred the highest or second highest predictive performance on all traits tested (**Fig.** 2), consistently exceeding the performance of the VEM-based `VIPRS`. At the same time, the main drawback of the grid search approach is that, despite the parallel software implementation, it results in a significant slowdown compared to the VEM approach (**Fig.** 3). In terms of predictive performance, the advantage of the grid search is most prominent for highly heritable traits, such as standing height and HDL (**Fig.** 2(a)). For the other traits, `SBayesR` is on-par or only marginally better. This indicates that the gap in predictive performance between `SBayesR` and the basic `VIPRS` model is mostly due to differences in hyperparameter estimation strategy.

**Figure 3:**
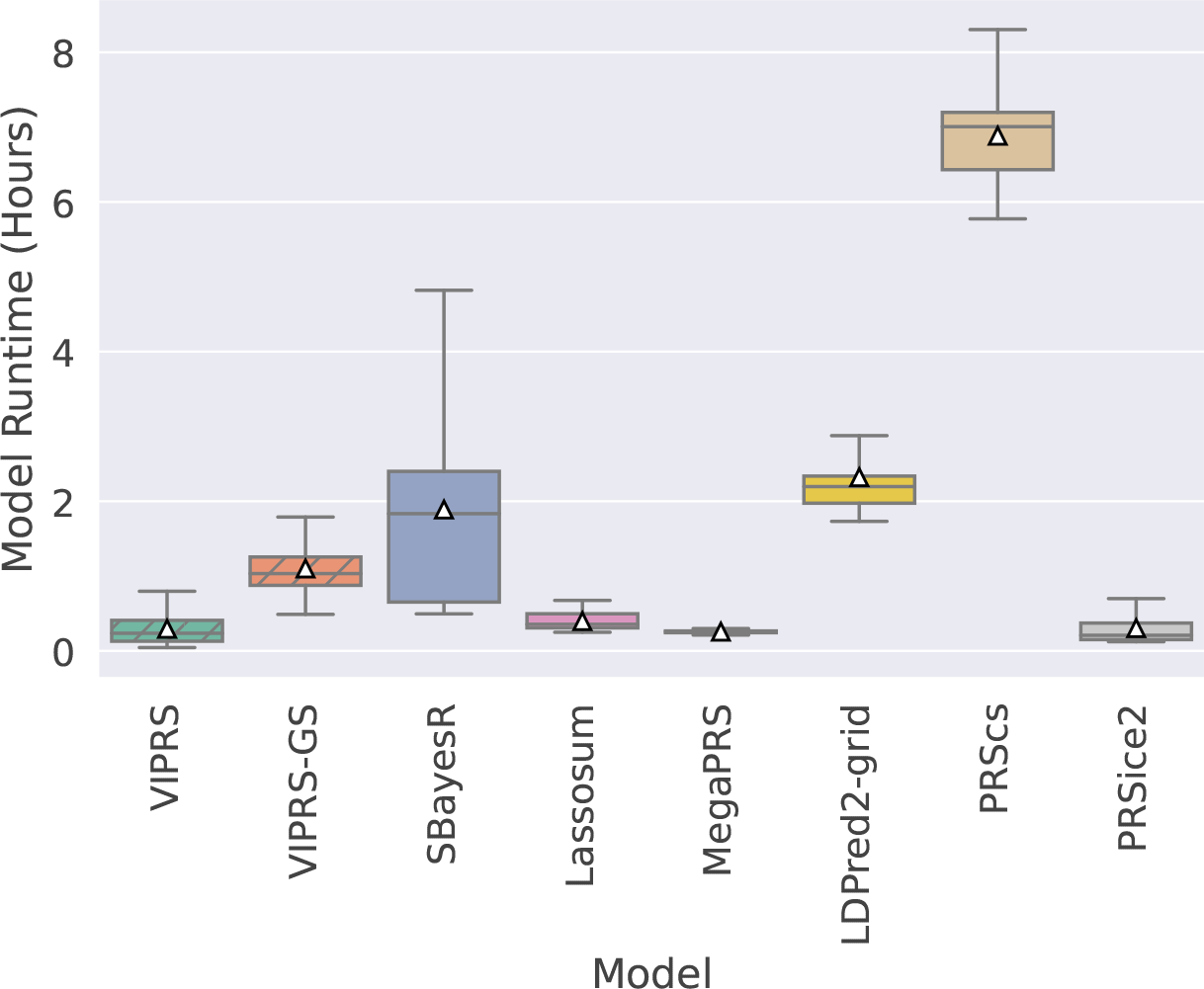
The total runtime (in hours) of the summary statistics-based PRS methods included in the study. The boxplot for each method shows the quartiles of the runtime from a total of 420 independent experiments (360 simulated traits plus 60 experiments on real measured traits, comprising the 12 phenotypes analyzed multiplied by the 5 training folds). The white triangles indicates the mean runtime for each method. The PRS methods shown are our proposed `VIPRS` and `VIPRS-GS` (using grid search to tune model hyperparameters) as well as 5 other baseline models: `SBayesR`, `Lassosum`, `MegaPRS`, `LDPred2` (grid), `PRScs`, and `PRSice2` (C+T). Dashed lines highlight the models contributed in this work.

### 2.4 PRS validation in minority populations in the UK Biobank

When trained on GWAS data from a single source population, transferability of PRS estimates across populations is limited [66, 67], with the degradation in prediction accuracy increasing with the increase in allele frequency differentiation (Fst) between populations [67]. At the same time, recent studies of cross-population genetic correlations have demonstrated strong correlations in the genetic architectures of complex traits between various ancestry groups [73, 74]. These correlations imply that PRS models that perform better in the source population will also tend to perform more favorably when applied to the target populations.

To assess this, we extracted genotype and phenotype data for individuals who self-identified as Italian (*N* = 6177), Indian (*N* = 6011), Chinese (*N* = 1769), and Nigerian (*N* = 3825). The self-reported ethnic backgrounds were further validated based on the Principal Components (PC) of the genetic relationship matrix (GRM) [67] (**Methods**). Using the effect size estimates derived from training the PRS models on summary statistics from the White British cohort across the five training folds, we computed a PRS for each individual in the target population. Given the real phenotype measurements for these individuals, we evaluated the predictive performance using relative incremental prediction *R*^2^, where the *R*^2^ in the target population was divided by the *R*^2^ of the best performing model on the test set in the White British cohort.

Our results confirm that for most of the traits analyzed, the models with the best predictive performance on the source population (White British) tend to transfer better to the target populations (**Fig.** 4). Furthermore, consistent with other analyses in this space [66, 67], the drop in prediction accuracy generally tracks with the Euclidean distance between the White British and the target populations in PC space. Interestingly, deviations from this general pattern were observed for LDL and birth weight, which may be due to gene-by-environment interactions [74]. For LDL specifically, we observed strong differentiation in transferability between PRS methods, with models employing variational inference techniques, `VIPRS` and `MegaPRS`, attaining upwards of 1.5 times the prediction accuracy of the next competing PRS method in individuals of Nigerian and Chinese ancestry (**Fig.** 4). This likely indicates that the `VIPRS` and, to a lesser extent, `MegaPRS` models correctly inferred the effect sizes of causal variants that are shared across ancestries but were missed by the other methods.

**Figure 4:**
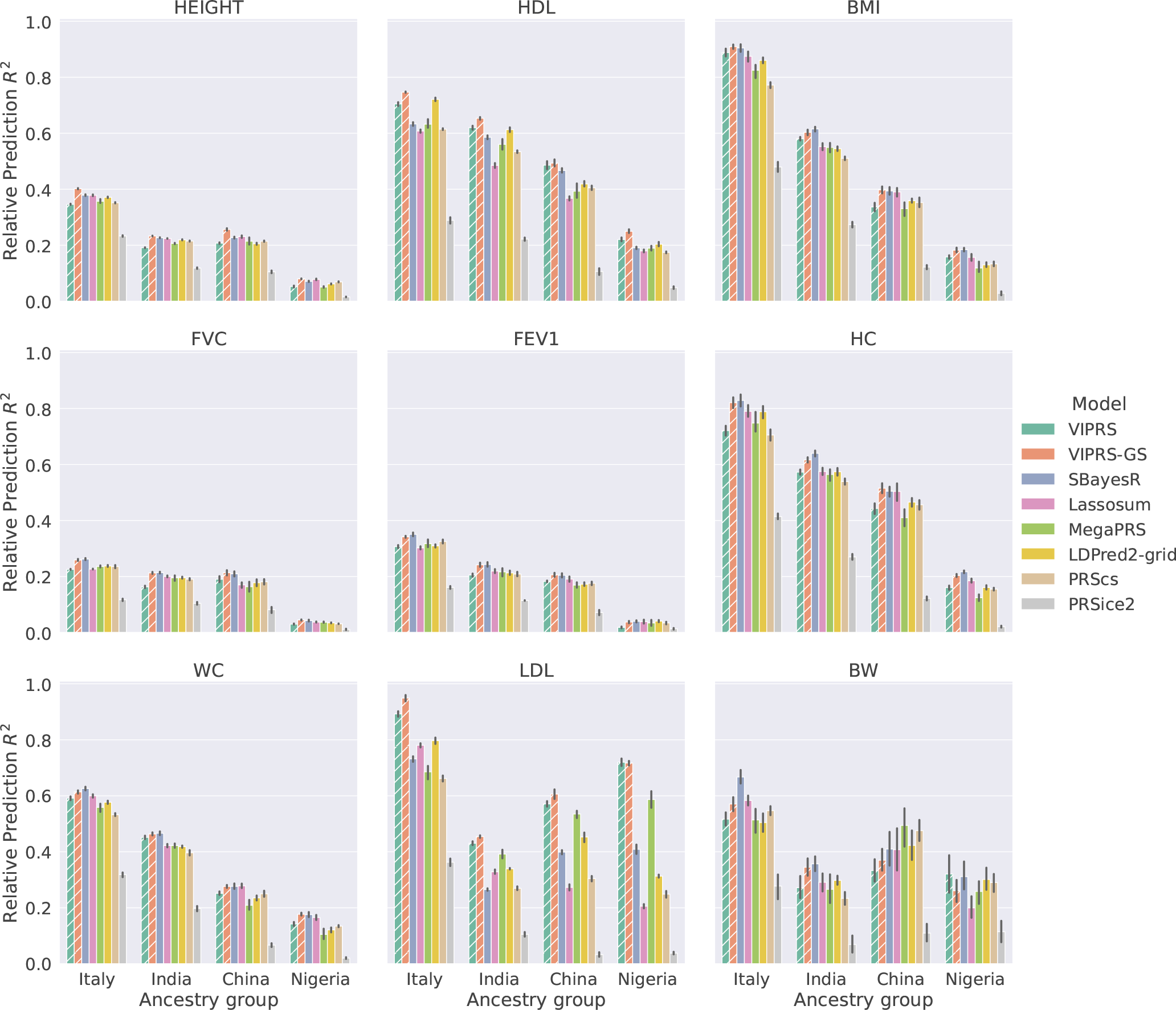
Relative predictive performance of summary statistics-based PRS methods on real quantitative phenotypes in minority populations in the UK Biobank. The PRS models were trained on summary statistics from the White British cohort in the UK Biobank using a 5-fold cross validation design. Then, the effect size estimates from the five training folds were used to perform predictions in individuals of Italian, Indian, Chinese, and Nigerian ancestry. Each panel shows the incremental prediction *R*^2^ in a given ancestry group relative to the prediction *R*^2^ of the best performing model on the White British cohort. The bars show the mean of the relative prediction metric across the 5 training folds and the black lines show the corresponding standard errors. The quantitative phenotypes analyzed are standing height (HEIGHT), high-density lipoprotein (HDL), body mass index (BMI), forced vital capacity (FVC), forced expiratory volume in 1 second (FEV1), hip circumference (HC), waist circumference (WC), low-density lipoprotein (LDL) and birth weight (BW). The PRS methods shown are our proposed `VIPRS` and `VIPRS-GS` (using grid search to tune model hyperparameters) as well as 5 other baseline models: `SBayesR`, `Lassosum`, `MegaPRS`, `LDPred2` (grid), `PRScs`, and `PRSice2` (C+T). Dashed lines highlight the models contributed in this work.

### 2.5 Scaling up VIPRS to 10 million variants

In recent years, with the advent of biobank-scale whole-genome sequencing efforts [3, 41] and improved variant imputation pipelines [33], there has been increasing interest in understanding the extent to which larger and larger sets of genetic variants enable us to better capture the genetic diversity underlying complex traits [75]. This is especially important in light of recent results that showed that a substantial proportion of the missing heritability is due to imperfect tagging of rare causal variants by common SNPs [75]. In this context, the main advantage of `VIPRS` is its speed and scalability (**Fig.** 3), thus we wanted to understand the extent to which our method could benefit from modeling an expanded set of SNPs. Here, following recent efforts in this space [32, 38], we examine the predictive performance of `VIPRS` with approximately 9.6 million measured or imputed genetic variants, almost an order of magnitude greater than the HapMap3 subset analyzed previously. This includes all bi-allelic variants with minor allele frequency greater than 0.1% and minor allele count greater than 5 in the White British cohort in the UK Biobank.

Following the same 5-fold cross-validation study design described earlier, we observed that modeling an expanded set of SNPs results in substantial improvements in prediction accuracy for highly heritable and polygenic traits, such as standing height and HDL, in comparison with the best performing PRS model using a subset of 1.1 million HapMap3 SNPs (**Fig.** 2), which is consistent with previous studies [6] (**Fig.** 5 and **Supplementary Fig.** S5). Concretely, for standing height and HDL in particular, including almost an order of magnitude more variants resulted in 3-7% relative improvement in the predictive performance of `VIPRS` and `VIPRS-GS`. However, the improvement is not consistent across all traits, especially for the VEM-based VIPRS. Similar to previous studies [6], we saw that, in some cases, including more variants led to modest drop in prediction accuracy, perhaps due to increased noise in the PRS estimate. Presumably, imputation errors for rare variants could potentially degrade the performance of the model in this setting. Therefore, we believe that this analysis presents a lower-bound on what could be achieved with more accurate and complete whole genome sequencing data [3, 41] (**Discussion**).

**Figure 5:**
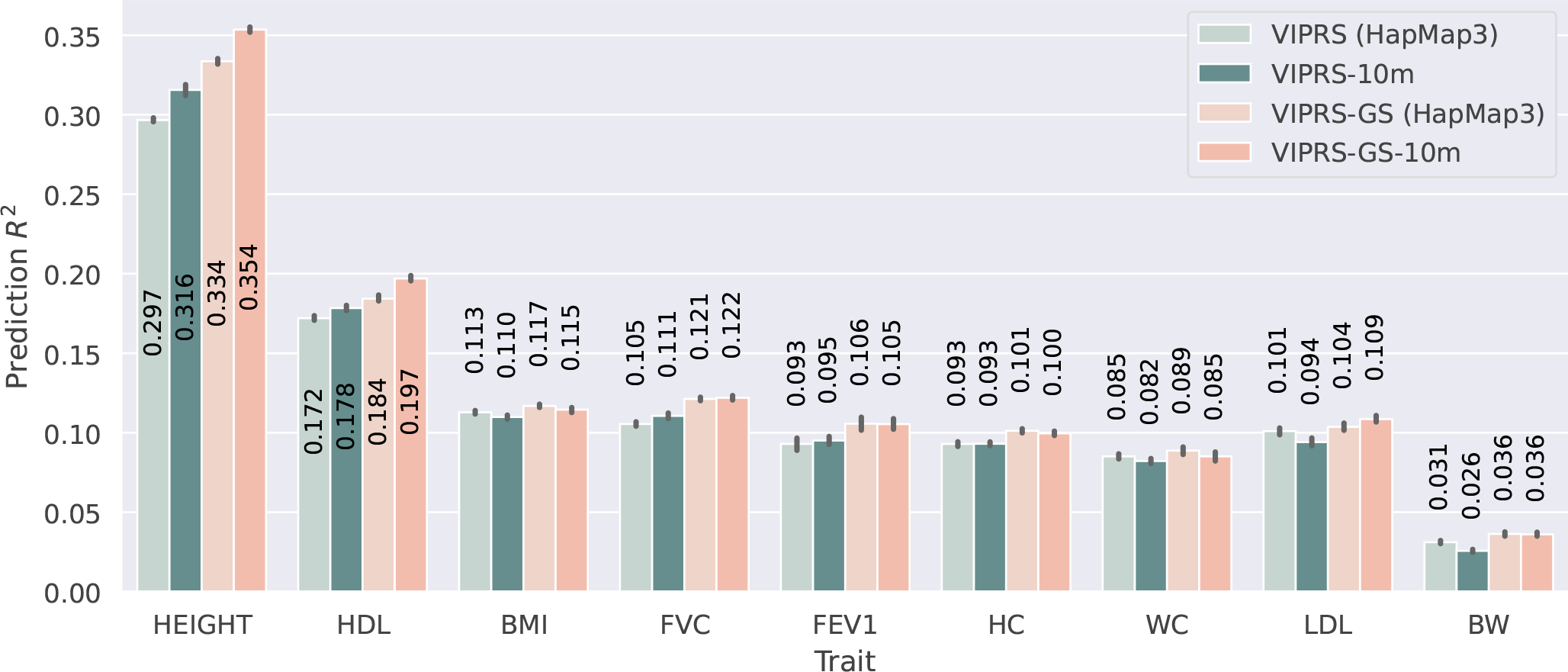
Comparing the predictive performance of the VIPRS method on real quantitative traits in the UK Biobank using the HapMap3 SNP set as well as an expanded set of 9.6 million genotyped and imputed variants. Each bar shows the predictive performance, in terms of incremental Prediction *R*^2^, of four different versions of the `VIPRS` model. From left to right, we have the standard `VIPRS` model trained on the HapMap3 subset (comprised of ≈ 1.1 million variants), `VIPRS-10m` is the `VIPRS` model trained on 10 million variants, `VIPRS-GS` is `VIPRS` with grid search trained on the HapMap3 subset, and finally `VIPRS-GS-10m` is the `VIPRS` with grid search trained on the 10 million SNPs. The black vertical lines show the standard errors across the five folds from the 5-fold cross-validation scheme. The quantitative phenotypes analyzed are standing height (HEIGHT), high-density lipoprotein (HDL), body mass index (BMI), forced vital capacity (FVC), forced expiratory volume in 1 second (FEV1), hip circumference (HC), waist circumference (WC), low-density lipoprotein (LDL) and birth weight (BW).

### 2.6 PRS analysis with external GWAS summary statistics

A common use case in the inference of polygenic scores involves settings where the GWAS summary statistics and the LD reference panel are estimated from two different cohorts [8, 35]. In other cases, the GWAS summary statistics may be derived from a meta-analysis that combines data from a number of different studies. These settings may present potential mismatches and heterogeneities between of LD reference panel and GWAS summary statistics and are thus challenging to model, often leading to substantial loss in predictive power [30, 31, 76, 77].

To systematically assess the robustness of `VIPRS` to potential heterogeneities and mismatches between the GWAS cohort and the LD reference panel, we conducted an analysis where we downloaded a number of publicly available GWAS summary statistics (PASS) for some of the traits analyzed previously, including: standing height [78], HDL and LDL [79], BMI [80], Type 2 diabetes [81], and Rheumatoid arthritis [82] (**Table** 1, see **Data Availability**). Most of these studies were conducted in individuals of general European ancestry, some in the form of meta-analysis. Therefore, we would expect some degree of differences between the LD reference panels derived exclusively from the White British cohort in the UK Biobank and the in-sample LD from these GWAS cohorts (which are not available).

We fit `VIPRS` as well as other PRS methods to the external GWAS summary statistics, providing the same 5-fold validation and testing cohorts within the UK Biobank for the purposes of hyperparameter tuning and evaluation as in the previous analysis. Our results indicate that, for some of the studies analyzed, the standard `VIPRS` model can be sensitive to mismatches between the GWAS cohort and LD reference panel (**Fig.** 6), though not to the same extent as `SBayesR`, which failed to converge for all of the quantitative traits analyzed, consistent with earlier work in this area [31]. Moreover, when using a validation set to tune the hyperparameters of the model, `VIPRS-GS` recovered most of the drop in performance relative to other PRS methods and showed competitive predictive ability (**Fig.** 6). This suggests that the `VIPRS` model trained to maximize the Evidence Lower BOund (ELBO) (**Methods**) of the external GWAS data may not generalize well to the UK Biobank individuals. Indeed, our experiments show a partial reversal in the correspondence between the training ELBO and the validation *R*^2^ (the metric that `VIPRS-GS` is optimizing) in the analysis of some of the external summary statistics (**Supplementary Fig.** S13), which explains the poor predictive performance of the basic `VIPRS` model in those settings.

**Figure 6:**
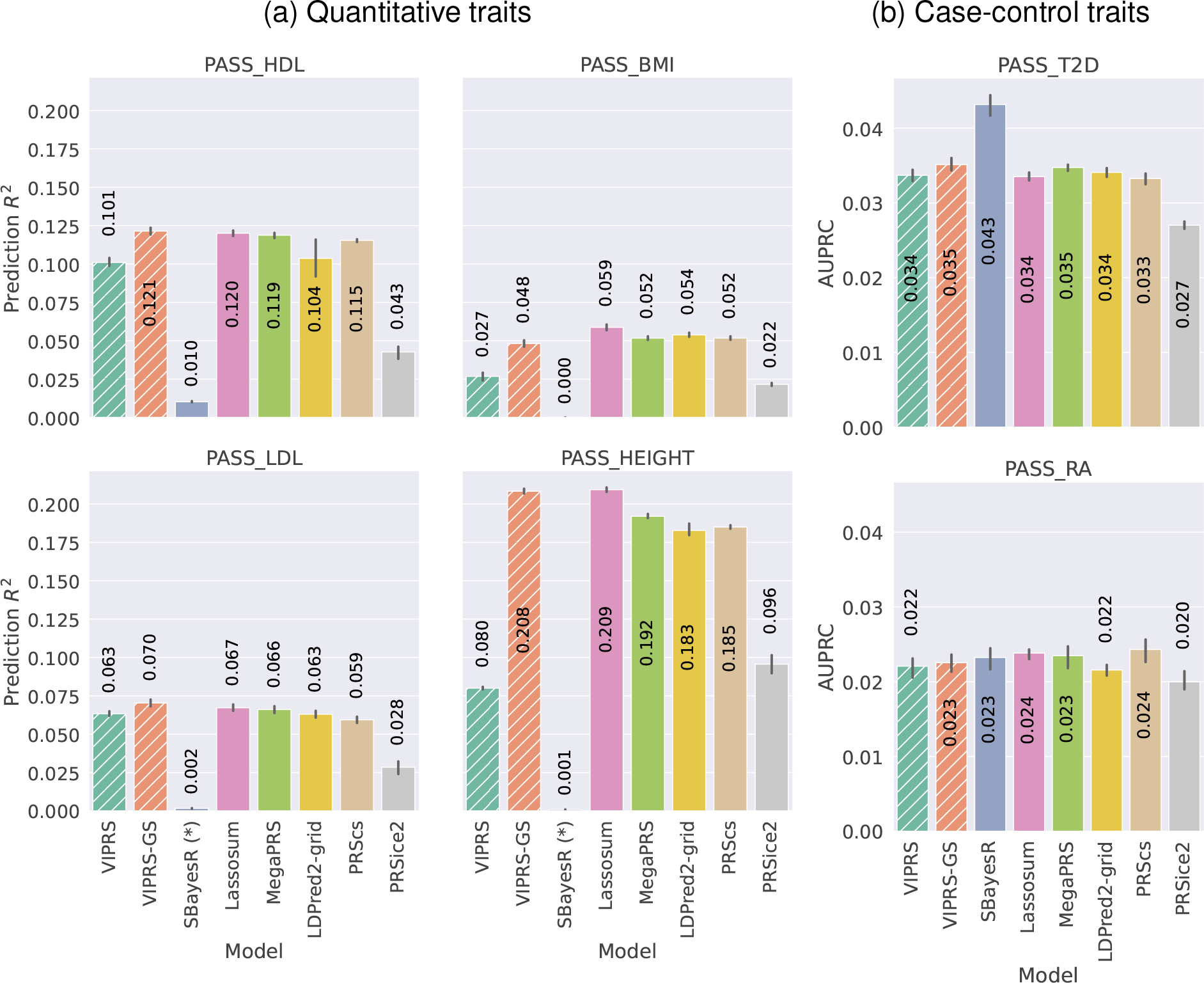
Predictive performance of summary statistics-based PRS methods on real **(a)** quantitative and **(b)** case-control phenotypes using external, publicly available GWAS summary statistics (PASS). Each panel shows the predictive performance, in terms of **(a)** incremental *R*^2^ and **(b)** area under the Precision Recall curve (AUPRC), of various PRS methods when applied to an independent test cohort in the UK Biobank. The bars show the mean and standard error of the prediction metrics across the 5 folds and the black lines show the corresponding standard errors. The quantitative phenotypes analyzed are standing height (PASS_HEIGHT), high-density lipoprotein (PASS_HDL), low-density lipoprotein (PASS_LDL). The binary phenotypes analyzed are type 2 diabetes (PASS_T2D) and Rheumatoid arthritis (PASS_RA). The PRS methods shown are our proposed `VIPRS` and `VIPRS-GS` (using grid search to tune model hyperparameters) as well as 5 other baseline models: `SBayesR`, `Lassosum`, `MegaPRS`, `LDPred2` (grid), `PRScs`, and `PRSice2` (C+T). The asterisk (*) next to the `SBayesR` method in panel **(a)** is to indicate that it did not converge on those traits. Dashed lines highlight the models contributed in this work.

Given this observation, if an independent validation set is not available, we recommend that users of the `VIPRS` software run principled tests of LD mismatch and heterogeneity, such as the recently published `DENTIST` method [77] before fitting the model to GWAS data. In the **Supplementary Information**, we also derived a stochastic estimator of the `DENTIST` test statistic that can be computed efficiently and provided that as a utility function in our software (see **Code Availability**).

## 3 Discussion

In this paper, we introduced `VIPRS`, a fast and flexible Bayesian PRS method that approximates the posterior for the effect sizes of genetic variants on the phenotype using variational inference techniques. Our genome-wide simulation analyses using genotype data from the White British cohort in the UK Biobank demonstrated that variational approximations to the posterior are not only computationally efficient, but they provide highly accurate polygenic score estimates across diverse genetic architectures. Indeed, in some simulation scenarios, `VIPRS` exceeded the predictive performance of competing Bayesian and non-Bayesian methods by large margins. The competitive prediction accuracy of the `VIPRS` method replicated in our analyses of real quantitative and binary phenotypes measured for the same UKB participants, though the differences between the methods in this setup were more modest. Similar systematic but mostly modest benefits were observed when PRS methods were applied to individuals from ancestries not included in the training dataset, emphasizing the robustness of the approach. For example, the effect size estimates by `VIPRS` for LDL cholesterol showed a large enough improvement in performance across ancestries to have potential clinical relevance [83] and make a significant dent in the transferability problem for that trait.

As highlighted throughout the text, we found that many implementation and modeling choices can have a substantial impact on the performance of the `VIPRS` model in analyses with GWAS summary statistics for real measured traits: hyperparameter tuning strategies, LD estimators, and the prior on the effect size all influenced the predictive performance in ways that varied across phenotypes and experimental setups. Overall, in most the setups and experimental conditions that we tested, the grid search approach for hyperparameter tuning combined with the spike-and-slab prior and windowed estimator of LD reliably outperformed or rivaled all the other variations of the model as well as previously described PRS methods. Notably, many of the individual modeling choices underpinning the `VIPRS-GS` model have been tried and tested in at least one other publication. Even the variational approximation that we derive bears some similarities to some existing methods that we compare against in our experiments, e.g. `MegaPRS` [32] (**Supplementary Information**). However, crucial details in the variational algorithm and its implementation and how they are joined together can still have significant impact on the overall performance, as illustrated by our experimental results.

One of the main strengths of the `VIPRS` model is its computational efficiency, which we exploited to test the predictive performance of the model with approximately 10 million SNPs, almost an order of magnitude greater than the standard HapMap3 subset used to train PRS methods [4, 6, 26, 30]. At this finer scale, we showed that modeling an expanded set of variants results in significant improvements in prediction accuracy for highly polygenic traits, such as standing height and HDL. This is consistent with recent whole-genome sequencing analyses which showed that a considerable proportion of rare causal variants are not well tagged by common SNPs [75]. There are a number of reasons that lead us to believe that the performance metrics that we report here are a lower bound on what could be achieved in modeling large-scale SNP array data. First, the vast majority of the variants that we added beyond the HapMap3 subset are rare and statistically imputed. Rare variant imputation is still a challenging problem and existing algorithms are known to have elevated error rates [84, 85]. We expect that these imputation errors can introduce substantial noise into the PRS estimate, and thus result in decreased prediction accuracy, as we observed for a number of the traits that we analyzed. This difficulty can potentially be addressed by using whole-genome sequencing (WGS) data for GWAS, which may soon be enabled by recent large-scale initiatives by the UKB [41] and TOPMed [3]. Second, residual confounding due to population structure may affect effect size estimation for rare variants [75, 86, 87]. In our GWAS pipeline, we corrected for population stratification using only the top 10 Principal Components (PCs) of the genetic relationship matrix (GRM), which may not adequately capture the more recent demographic history reflected by rare variants [87, 88]. This residual confounding effect may be addressed by increasing the number PCs used in the GWAS analysis [75] or utilizing more genealogically-informed estimates of the GRM [89].

Despite its competitive predictive ability, we believe that there are a number of modeling choices underlying `VIPRS` that can potentially be improved in future work. Firstly, compared to simulated phenotypes, the generative process for real traits is unknown and likely involves complex and heterogeneous genetic architectures that are not well described by a two-component Gaussian mixture prior. The spike-and-slab prior assumes that all genetic variants have a uniform prior probability of being causal and that the causal SNPs have equal expected contribution to the heritability, which is a simplistic assumption given what is known about the genetic architectures of complex traits [12–14]. This motivated us to explore a more general and flexible Gaussian mixture prior with four mixture components [6, 24]. Our experimental results show that adding mixture components improves accuracy for highly heritable and polygenic traits, such as standing height, but did not systematically improve accuracy for less heritable traits, perhaps because of reduced power to identify the larger number of parameters. Future work using priors informed by functional annotations (e.g. [32, 90]) is a promising avenue to improve accuracy in these cases. Second, our validation analyses in the UKB confirmed that, in general, `VIPRS` as well as other PRS methods do not transfer well across populations or ancestry groups, despite some notable differences between the methods. Recent work has highlighted that transferability in the context of summary statistics-based PRS methods is best achieved when we jointly model the effect sizes of multiple ancestrally homogeneous populations within the same framework [55, 91, 92]. This formulation has proved successful for some Bayesian PRS methods [55, 92] and we believe that fast variational approximations to the posterior under such models will increasingly be shown to be effective and highly competitive.

Finally, while our results showed that variational approximations to the posterior are a promising alternative to MCMC techniques in predictive settings, it is important to highlight that meanfield variational approaches are known to underestimate the posterior variances and covariances in some cases [43, 93, 94]. In practice, this means that the variational posterior will tend to underestimate the proportion of variance explained (e.g. **Supplementary Fig.**S8 and S9) [48] and may also result in miscalibrated PRS confidence intervals, if such a quantity is sought for some downstream applications [36]. This limitation can be addressed with more expressive variational families [95], such as those derived with variational boosting [96], or alternatively, with help of modern Bayesian inference techniques that combine variational methods and MCMC [97].

## 4 Methods

### 4.1 Detailed description of the VIPRS model

We formulate the problem of polygenic prediction in terms of the standard linear model,

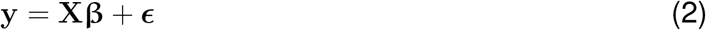

where **y** is an *N* × 1 vector of phenotypic measurements for *N* individuals, **X** is the *N* × *M* genotype matrix that records the counts of alternative alleles for each individual at each genetic marker, **β** is a vector of effect sizes for each of the *M* markers, and ***ϵ*** is an *N* × 1 vector that captures the residual effects on the trait for each individual. For quantitative traits, we assume that the phenotypes follow a Gaussian likelihood, such that 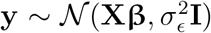, with 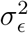 being the residual variance. Practically, a central goal of polygenic risk modeling is to arrive at a robust estimate for the effect sizes **β**. In the Bayesian framework, this problem is tackled by imposing a prior distribution over the effect sizes and then deriving a solution for the posterior distribution given the data likelihood and the prior,

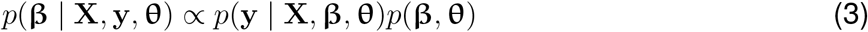

where **θ** encapsulates all fixed hyperparameters in the model, i.e. parameters that we do not assign a prior. Here, the constant of proportionality is the marginal likelihood or the partition function for the posterior, ∫ *p*(**y** | **X**, **β**, **θ**, **s**)*p*(**β**, **θ**)*d***β**, also known as the model evidence [39, 40]. In recent years, considerable work has been devoted to deriving Bayesian PRS models with flexible priors on the effect sizes, such as the continuous shrinkage [26] and mixture priors [6]. In this work, we follow the lead of earlier approaches, e.g. [4, 49], and assign a spike-and-slab prior [56, 57] on the effect sizes,

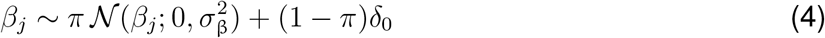

Here, *π* is a parameter that denotes the prior probability of a variant being causal, 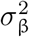 is the prior variance on the effect size of each SNP, and *δ*_0_ is the Dirac delta function. In the simplest formulation of this model, we assume that *π* and 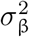 are shared across all SNPs. Thus, *π* may also be considered as the fraction of variants that are causal for the trait of interest, and the 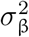 parameter is related to the trait’s per-SNP heritability [4, 68]. The spike-and-slab prior is a special case of the more general mixture prior:

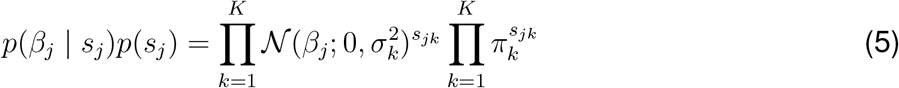

where *s_jk_* is binary indicator for SNP *j* belonging to the *k^th^* mixture component, with *π_k_* and 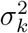 denoting the the mixing proportion and prior variance for component *k*, respectively. It is well known that Bayesian linear regression models with a spike-and-slab prior on the effect sizes result in an intractable posterior [49, 56, 57], necessitating the use of approximate posterior inference schemes.

In most of the previous Bayesian PRS formulations, the authors employ a Gibbs sampler, a Markov Chain Monte Carlo (MCMC) technique that relies on conditional conjugacy between the prior and the likelihood, to approximate the posterior distribution of the effect sizes [4, 6, 26, 30]. In this work, we instead leverage a technique known as Variational Inference (VI) [42], which approximates intractable densities by proposing a simple parametric distribution *q*(**β**, **s**) and optimizing its parameters to match the true posterior as closely as possible [43]. The closeness between the true posterior and the proposed distribution is measured by the Kullback-Leibler (KL) divergence,

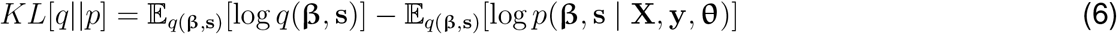

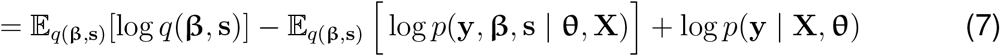

where 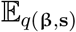 is the expectation taken with respect to the proposed distribution [39, 40]. However, the KL divergence includes the normalizing constant that made the posterior intractable in the first place. Thus, practitioners typically optimize a surrogate objective known as the Evidence Lower BOund (ELBO) of the log marginal likelihood [39, 40, 43]:

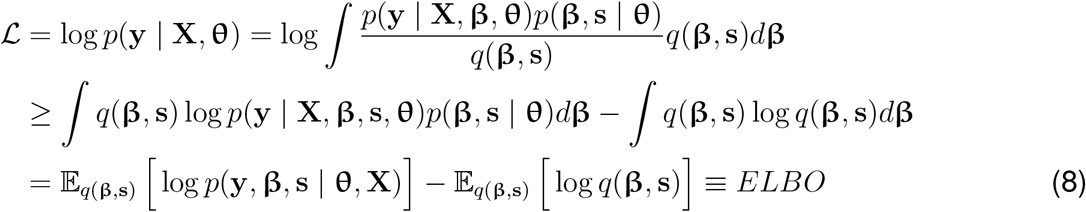

Here, the first term in (8) is the expectation of the log joint likelihood of the phenotypes and the effect sizes and the second term corresponds to the negative entropy of the variational distribution. The ELBO (8) and the KL-Divergence in (7) add up to the marginal likelihood: 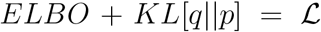. Therefore, maximizing ELBO is equivalent to minimizing the KL-Divergence [39, 40, 43].

The choice of approximating variational distribution *q*(**β**, **s**) is a central component in this setting. For simplicity and computational efficiency, we make use of the paired mean-field assumption [39, 40, 59], whereby the density factorizes across the input coordinates, and model the effect size at each locus with a two component Gaussian mixture density [49, 59, 72],

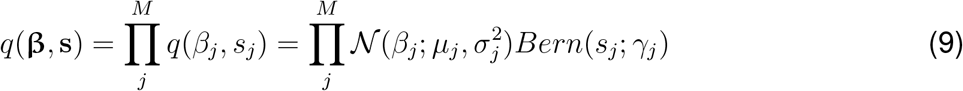

Here, *μ_j_*, 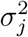, *γ_j_* are the variational parameters defined for each variant in the dataset and 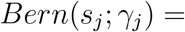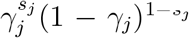 denotes a Bernoulli distribution with probability *γ_j_* for SNP *j*. Therefore, the Bernoulli indicator in the proposed distribution approximates the posterior probability that the variant is causal for the trait of interest and the Gaussian component approximates the posterior for the effect size [49]. In the **Supplementary Information**, we provide detailed derivations which show that, under certain assumptions, this variational family leads to the following closed form updates that only depend on GWAS summary statistics and the SNP-by-SNP correlation or Linkage Disequilibrium (LD) matrix, which can be derived from an appropriately-matched reference panel:

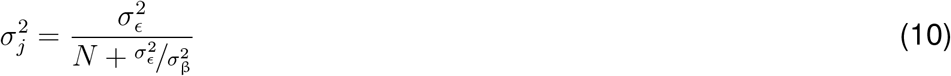

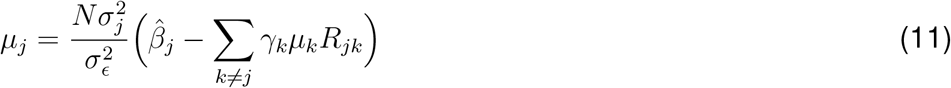

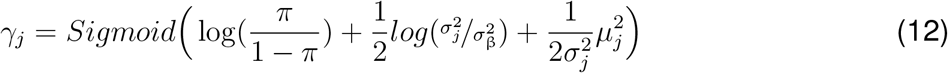

Here, 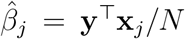 is the standardized marginal GWAS effect size and ***R*** is the LD matrix. This formulation enables us to employ a fast coordinate ascent algorithm to approximate the posterior distributions of the effect sizes.

In addition, to perform inference given some of the unknown fixed parameters, i.e. 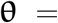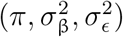, the basic formulation of the `VIPRS` model uses the Variational Expectation-Maximization (VEM) algorithm [48, 49, 72], where, in an alternating fashion, in the E-Step we update the variational parameters given the hyperparameters and in the M-Step we update the hyperparameters of the model. In both the E- and M-steps, we update the free parameters of the model to maximize our objective, the ELBO. In **Supplementary Information**, we show that updating the hyperparameters to maximize the ELBO also results in closed-form solutions for those parameters.

Despite the conceptual simplicity of the model described above, fitting such a model to biobank-scale data presents several computational challenges. For instance, the closed-form update equations for some of the variational parameters involve terms that relate to the LD between the focal variant and all other variants in the genome. This can be computationally prohibitive to compute for millions of variants and for hundreds of EM iterations. To overcome this, we follow the lead of other summary-statistics-based PRS methods and use a banded or shrunk LD matrix [4, 6, 26, 30], which results in substantial improvements in speed without substantially degrading predictive performance.

### 4.2 Hyperparameter tuning strategies

The standard VIPRS model employs a Variational EM framework to infer the hyperparameters 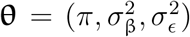, where in the M-step, we update each hyperparameter to maximize the surrogate objective, i.e. the ELBO [49, 72]. This strategy works well in many settings, but it is prone to entrapment in local optima [72], which may degrade overall predictive performance of the model. In this work, we explored three alternative strategies for tuning the hyperparameters of the model.

#### VIPRS-GS

In the first strategy, we performed grid search (GS) over the hyperparameters of the model, selecting the values that result in the best predictive performance on a held-out validation set [25, 27, 30]. We also explored a pseudo-validation variant of the model (**Supplementary Information**) and showed that it results in almost identical prediction accuracy (**Supplementary Fig.** S6 and S7). The grid search was performed specifically over the proportion of causal variants *π*, with the remaining parameters updated according to their maximum likelihood estimates. The grid for *π* ranged from 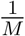 to 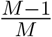 with 30 equidistant values on a log_10_ scale, where *M* is the number of variants included in the model.

#### VIPRS-BO

To search over the hyperparameter space without the constraint of a predefined and discrete grid, we experimented with a second hyperparameter tuning technique known as Bayesian Optimization (BO) [62]. In Bayesian Optimization, we assume that there is an underlying unknown function *f*(**θ**) that takes the hyperparameters as input and outputs a certain score that we wish to optimize, such as the training ELBO or the validation *R*^2^. This unknown function is modeled with a Gaussian Process (GP) prior, which allows us to explore the parameter space efficiently while accounting for uncertainty in a principled manner. The other component in this framework is the acquisition function, a heuristic that maps from the GP posterior to information about the most promising regions in hyperparameter space [62, 98]. In our experiments, we used the `scikit-optimize` python package to perform this optimization, with `gp_hedge` as the default acquisition function. The optimizer was allowed to sequentially evaluate up to 20 points in a bounded 1*D* space for the hyperparameter *π*.

#### VIPRS-BMA

In the third strategy, we used a Bayesian Model Averaging (BMA) framework, where we use importance sampling to integrate out some of the hyperparameters of the model, as outlined in [49]. The main idea here is that instead of fixing *π* to a particular value, we fit the `VIPRS` model along a grid of *π* values, as in `VIPRS-GS`, and then take a weighted average of the effect size estimates for each SNP based on each model’s ELBO [48, 49].

Similar to previous work in this area, we note that these three strategies can be deployed in conjunction with the VEM framework [48, 49, 99], where some of the hyperparameters are updated using their maximum likelihood estimates while the remaining parameters are optimized via the user’s strategy of choice. This is important in practice, because with the three hyperparameters of the model, an exhaustive search will require searching over a three-dimensional grid, which can be computationally expensive. Therefore, in our experiments and analyses, for all of the three strategies that we explored, we only implemented a search over the fraction of causal variants *π* and estimated the other two hyperparameters using the closed-form updates in the M-Step.

### 4.3 Data preprocessing

To assess the performance of `VIPRS` on a biobank-scale dataset, we made use of the UK Biobank (UKB), a large database of genomic and phenotypic measurements from 488,377 participants from the United Kingdom [1]. In its latest release, the UKB database has genotype information from 488,377 individuals, from which, after applying standard quality control procedures, we retained data for a total of 337,205 samples. Briefly, the sample quality controls involved selecting unrelated individuals with White British ancestry, defined by the UKB based on self-reported ethnic background as well as Principal Components Analysis (PCA) of the GRM, and who were also included in the PCA and phasing procedures outlined in [1]. We restricted our main analyses to the White British cohort in order to maximize power to detect causal effects while reducing confounding. In addition this, we filtered data for individuals with detected sex chromosome aneuploidy, excess relatedness, or missing genotype rate exceeding 5% from this analysis.

The genetic variants or SNPs included in the study were selected based on a number quality control filters, applied at various stages in the analysis. For the base dataset, we excluded variants with duplicate `rsIDs`, ambiguous strand, imputation quality score < 0.3, Hardy-Weinberg Equilibrium p-value < 10^−10^, or genotype missingness rate > 0.05. We also removed multiallelic variants as well as SNPs in long-range LD regions, as specified in Supplementary Table 13 in Bycroft et al. 2018 [1]. In the GWAS analyses or LD matrix construction, we further filtered variants with minor allele count (MAC) < 5 or minor allele frequency (MAF) < 0.1%. This resulted in a total of 9,590,026 bi-allelic variants that were used in the expanded SNP set analyses. Finally, following standard practice in PRS methodologies [6, 30], for the base analyses with the `VIPRS` model, we restricted to the set of variants in the HapMap3 reference panel [34], resulting in a total of 1,093,308 SNPs. Most of these quality control procedures were carried out with the genetic analysis software tool `plink2` [100].

### 4.4 Construction and efficient representation of LD matrices

An important quantity in the model is the Linkage Disequilibrium (LD) or SNP-by-SNP correlation matrix ***R***. The matrix, or its columns, show up mainly in the update equations for the variational parameters *μ_j_* of each SNP *j* (11), the estimate of the residual variance 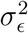, as well as in the objective function (i.e., ELBO) (**Supplementary Information**). Our software supports a number of different LD matrix estimators, including sample, block, shrinkage, and windowed estimators.

#### Sample estimator

In the sample estimator, we estimate the sample Pearson correlation coefficient between all SNPs on the same chromosome, which results in a dense matrix. For larger chromosomes and SNP sets, it is impractical to load dense matrices of this scale to memory. To handle data at that scale, we use compressed and chunked on-disk storage with `Zarr` arrays in `python` for fast, multi-threaded read and write access. Then, as we iterate through SNPs in the E-Step, we load the matrix into memory one-chunk at a time, thus allowing us to train `VIPRS` with extremely large LD matrices. In **Supplementary Information**, we describe a procedure that allows us to load the LD matrix only once per iteration, resulting in improved speed and efficiency.

#### Block estimator

In the block LD estimator, we only estimate the sample LD between SNPs that are within the same LD block, as defined by, e.g. LDetect [70]. This is similar to what is done in the `Lassosum` and `PRScs` frameworks [25, 26].

#### Shrinkage estimator

In the shrinkage estimator, we shrink and threshold the entries of the sample LD matrix according to procedure outlined by [6, 69] and implemented in the `gctb` software. Briefly, for the shrinkage estimator, we shrink each element of the LD matrix by a quantity proportional to the distance between pairs of variants *j* and *k* in along the chromosome: 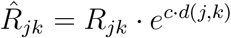. In this context, *d*(*j, k*) is the distance in centi Morgans between variants *j* and *k* and the constant *c* is related to sample size used to infer the genetic map as well as effective population size [6, 69].

#### Windowed estimator

For the windowed LD estimator, we only consider the correlation between a focal variant with variants that are at most 3 centi Morgan away from it along the chromosome [30, 68]. This estimator results in compact and banded LD matrices that can easily fit in memory on modern compute nodes.

To construct LD matrices for the main analyses of this paper, we selected a random subset of 50,000 individuals from the White British cohort described above. Within that group of individuals, we filtered SNPs with minor allele count (MAC) < 5 or minor allele frequency (MAF) < 0.1%, and again restricted to variants in the HapMap3 reference panel. For the analyses with the expanded set of variants, we only removed the HapMap3 filter. Unless explicitly stated otherwise, the analyses with the `VIPRS` model employed the windowed estimator for LD, with the distnace cutoff set to 3cM. The matrices are stored in compressed `Zarr` array format and are publicly available for download (see **Data Availability**).

### 4.5 Simulation study

To assess the predictive performance of `VIPRS` on large-scale datasets and for varying genetic architectures, we conducted a genome-wide simulation study using the pre-processed genotype data from the UK Biobank cohort. We simulated quantitative and binary traits according to six different genetic architectures and three settings for the additive genetic variance, 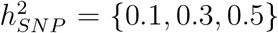, for a total of 18 simulation configurations for each trait category (continuous and case-control). For the first three genetic architectures, we simulated the effect size for each variant according to the generative model outlined previously in Equation 4, with three settings for the proportion of causal variants, *π* = {10^−4^, 10^−3^, 10^−2^}. The next two simulation scenarios involved sampling the effect size for each variant from a scale mixture of Gaussians (**Supplementary Information**), 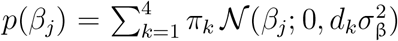, with the mixing proportions set to *π* = {0.95, 0.02, 0.02, 0.01}. The variance multipliers *d_k_* were set to *d* = {0.0, 0.01, 0.1, 1} for the sparse mixture model and *d* = {0.001, 0.01, 0.1, 1} for the infinitesimal mixture model. Finally, the last genetic architecture tested was the standard infinitesimal model, with the effect size drawn from a zero-centered Gaussian density 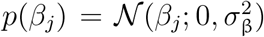. For each configuration, we generated 10 independent phenotypes, for a total of 180 simulated traits. For the binary traits, we followed the same procedure, but used the liability threshold model [63] to obtain case-control status, with prevalence set to 15%.

After we generated simulated phenotypes for all individuals in the study (*N* = 337, 205), we excluded the 50,000 samples used to generate the LD matrices and randomly split the remaining samples into 70% training (*N* = 201, 043), 15% validation, and 15% testing (*N* = 43, 081 each). The genotype and simulated phenotype data of the training samples were then used to generate GWAS summary statistics using `plink2` [100].

### 4.6 Application to real traits from the UKB

To assess the predictive performance of `VIPRS` on real phenotypes, we extracted phenotypic measurements for 9 quantitative and 3 case-control traits for the UKB cohort described previously. The quantitative phenotypes included log-transformed Waist Circumference (WC), log-transformed Hip Circumference (HC), Standing Height (HEIGHT), Birth Weight (BW), logtransformed Body Mass Index (BMI), log-transformed High-Density Lipoprotein (HDL), Low-Density Lipoprotein (LDL), Forced Vital Capacity (FVC), and Forced Expiratory Volume in the first second (FEV1). For each trait, we excluded samples with outlier or extreme values for the trait. For the remaining samples, within each sex separately, we corrected for age and the top 10 Principal Components (PCs) of the genetic relationship matrix (GRM), and then applied a Rank-based Inverse Normal Transform (RINT) on the residuals [101]. To assess the predictive performance on held-out test sets, we performed 5-fold cross-validation. For each split, the training data was further split into 90% training and 10% validation to facilitate running PRS methods that require a validation set to tune their hyperparameters.

The case-control phenotypes included in the analysis are Asthma (prevalence 12.7%), Type 2 Diabetes (T2D) (prevalence 2.3%), and Rheumatoid arthritis (RA) (prevalence 1.7%). To assess the predictive performance on held-out test sets, we performed stratified 5-fold cross validation, followed by splitting the training data into 90% training and 10% validation in a stratified manner to keep the prevalence approximately the same for all subsets of the data.

The phenotypes and associated sample sizes in the UK Biobank are listed in Table 1. The detailed scripts with the extraction and transformation procedure for each phenotype is included in the public repository associated with this publication (**Code Availability**). The 5-fold cross validation procedure was performed using the `scikit-learn` package in `python` [102].

### 4.7 Validation in minority populations in the UKB

To validate the relative predictive ability of `VIPRS` in individuals of different backgrounds, we used the approach of Prive et al. [67] to identify subgroups of relatively uniform ancestry and ethnicity. Using self-reported ethnic background as well as PCA medoids from [67], we extracted genotype data for individuals of Italian (*N* = 6177), Indian (*N* = 6011), Chinese (*N* = 1769), and Nigerian (*N* = 3825) ancestry. In genetic analyses, those ancestry groups show various levels of allele frequency differentiation (Fst) when compared to the White British cohort [67]. The samples were selected after applying the same quality control filters as before. Mainly, we retained individuals who were used in the PCA and phasing procedures and filtered samples with detected sex chromosome aneuploidy, or excess relatedness from this analysis.

For each individual in those target populations, we extracted phenotype data for the traits analyzed previously (**Table**1). Then, we used effect size estimates derived from the 5-fold analyses on the White British cohort to generate polygenic scores for individuals in those minority populations. Given these polygenic score estimates, we computed the relative prediction *R*^2^ as the incremental *R*^2^ in the target population divided by the *R*^2^ of the best performing PRS model on the test set in the White British cohort. This metric is designed to highlight the transferability of PRS estimates across different population and ancestry groups.

### 4.8 PRS method comparison

To compare the predictive performance of VIPRS to state-of-the-art models for polygenic risk prediction with summary statistics, we included a diverse collection of methods with different assumptions and implementations, including three stochastic Bayesian methods `SBayesR` (`gctb 2.03`) [6], `PRScs-auto` [26], and `LDPred2-grid` (`bigsnpr 1.9.11`) [30], a variational Bayesian PRS method `MegaPRS` (the `BayesR` variant with the `GCTA` heritability model as implemented in `LDAK 5.2`) [32], a penalized regression method (`Lassosum 0.4.5` [25]), and finally, as a simple baseline, we included a a C_+_T method (`PRSice2 2.3.5` [27]). In addition to being widely used in practice, most of these methods were selected because they have been shown to be competitively accurate in recent comprehensive surveys [37, 38, 64].

For each method, we provided the GWAS summary statistics for the simulated and real phenotypes and ran the model with default or recommended settings. For models that require a validation set to tune the hyperparameters, we provided a held-out validation subset of the data. Once the models converge, we output the effect size estimates and then generated polygenic scores for the samples in the test set. Given these polygenic scores, the models were then evaluated for the quality of their predictions.

For quantitative traits, we reported the incremental prediction *R*^2^, defined as the *R*^2^ of a linear model with the PRS and covariates (age, sex, and top 10 PCs) minus the *R*^2^ obtained from a linear model with the covariates alone. For case-control phenotypes, we reported the area under the Precision-Recall curve (AUPRC) between the polygenic score and the binary phenotype. In addition to these prediction metrics, we also compared the run-time (wall-clock time) of the different methods to gauge their scalability and computational efficiency.

The detailed specification of priors, grid values, and other hyperparameters for each PRS model is shown in the repository accompanying this manuscript (see **Code Availability**).

### 4.9 Software implementation

The data structures and inference algorithms for the `VIPRS` model are implemented in two `python` packages that are open source and publicly available on GitHub (see **Code Availability**). The first software package, `magenpy`, implements scripts and routines for computing LD matrices and transforming them to `Zarr` array format, simulating complex traits from genotype data, and harmonizing multiple genetic data sources, such as GWAS summary statistics, LD reference panels, functional annotations, etc. The second software package, `viprs`, implements the optimized variational inference algorithms to obtain posterior estimates for the effect sizes. For optimal speed and efficiency, the coordinate ascent routine is written in `cython`, a compiled programming language that produces `python`-compatible modules with minimal overhead [103]. Both software packages follow object-oriented design principles to allow for streamlined user extensions and experimentation by experienced developers. We also provide runner scripts that allow users to perform inference using commandline interfaces.

## Code Availability

- The code to process GWAS summary statistics, construct LD matrices, and provide necessary data structures for the `VIPRS` model is publicly available on GitHub: https://github.com/shz9/magenpy
- Code to run the `VIPRS` model and perform posterior inference is available via GitHub: https://github.com/shz9/viprs
- Scripts to replicate all the analyses and generate the figures for this manuscript are available via GitHub: https://github.com/shz9/viprs-paper

## Data Availability

- The external GWAS summary statistics were downloaded from the LD-Score regression repository: https://alkesgroup.broadinstitute.org/LDSCORE/all_sumstats/
- Pre-computed LD matrices in `Zarr` format are available for download via Zenodo: https://doi.org/10.5281/zenodo.7036625

## Acknowledgements

We thank members of the Li and Gravel labs for useful feedback and discussions on this project. We are particularly grateful to Doruk Cakmakci and Goodarz Koli Farhood for testing various aspects of the software. We also thank Hans Markus Munter, Stephen Sawcer, and Mark Lathrop for help with data access and Celia Greenwood and Mathieu Blanchette for feedback on the project at various stages. We are grateful to Doug Speed, Tianyuan Lu, Lino Fonseca Ferreira for feedback and discussion on earlier drafts of this manuscript. The UK Biobank analyses in this work were conducted under Application 4408. Y.L. is supported by Natural Sciences and Engineering Research Council (NSERC) Discovery Grant (RGPIN-2019-0621), Fonds de recherche Nature et technologies (FRQNT) New Career (NC-268592), and Canada First Research Excellence Fund Healthy Brains for Healthy Life (HBHL) initiative New Investigator start-up award (G249591).

## Author contributions

S. G. and Y. L. jointly supervised this work. Y.L. conceived the study. Y.L. and S.Z. developed the methodology. S.Z. implemented the computational software and performed all the experiments. S.Z., Y.L., and S.G. analyzed the results. S.Z. wrote the initial manuscript. All of the authors wrote the final version.

## Supplementary Information

### S1 VIPRS model derivation and implementation details

#### S1.1 Model formulation

We model the dependence of a quantitative phenotype *y* on the genotype matrix **X** via the standard linear model:

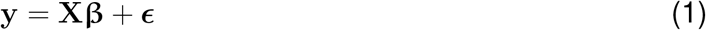

where **y** is a vector of phenotypic measurements for *N* individuals, **X**_*N*×*M*_ is a matrix of allelic counts for each individual and genetic marker, **β** is a vector of effect sizes, and ***ϵ*** is a vector of residual effects for each individual. We assume that the phenotype conditioned on the effect sizes and genotypes follows an isotropic Gaussian likelihood 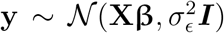. In Bayesian inference, we treat all unknown parameters as latent random variables and assign them a prior distribution. We assume that the effect sizes **β** have, as a prior, a factorized Gaussian mixture distribution with *K* components [6, 48],

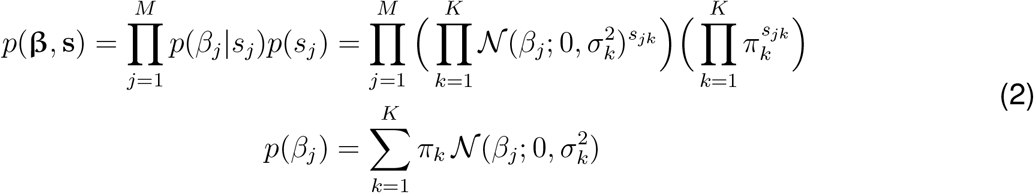

where *s_jk_* is a latent random variable that takes value 1 if SNP *j* belongs to component *k* in the mixture, and 0 otherwise. The parameters *π_k_* and 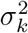 are the mixing proportions and the prior variance for component *k*, respectively. To allow for sparsity in the model structure, we assume that one of the components, e.g. the last component, is a degenerate Gaussian density with zero variance (i.e., the Dirac delta density *δ*_0_). In this case, when *K* = 2, the above mixture prior reduces to the standard spike-and-slab prior [56–58], where the spike corresponds to the null component and the slab density corresponds to the non-null component. Given this model structure, the goal of Bayesian inference is to estimate the posterior for the effect sizes,

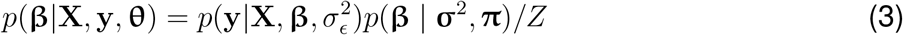

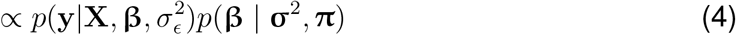

where 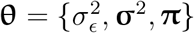 is a collection of hyperparameters for the model and 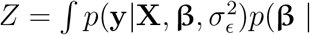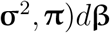 is the marginal likelihood or the partition function. Therefore, the posterior is proportional to the product of the likelihood and the prior.

#### S1.2 Variational inference

The mixture prior on **β** results in intractable posterior because the analytical closed-form of the marginal likelihood is not available. Thus, we need to devise a scheme for approximate posterior inference. In this work, we use variational inference, a deterministic approximate inference scheme that involves specifying a parametric distribution family *q*(**β**, **s**) and optimizing its parameters to match the true posterior as closely as possible [39, 40, 43]. We use the same family of distributions as the prior, which assumes that the effect size at each SNP follows a mixture of *K* Gaussians (with the last Gaussian as a spike with zero variance) [48]. For tractability and computational efficiency, we make use of the paired mean-field assumption [59], which factorizes the joint distribution of the effect sizes and causal indicators into a product of independent densities for each SNP,

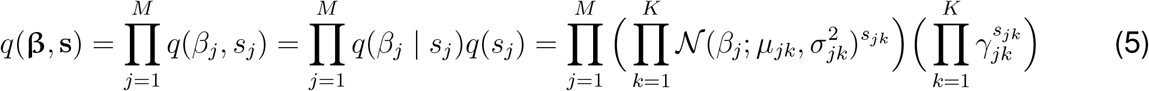

where *μ_jk_*, 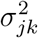, *γ_jk_* for all *j* and *k* are the variational parameters that require tuning. Given this posterior distribution family, for notational convenience, we define the first and second moments for the effect size of SNP *j*,

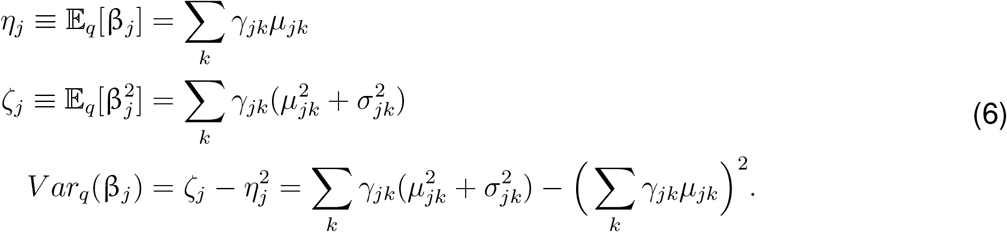

where 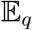 and *V ar_q_* denote the expectation and variance taken with respect to the proposed distribution *q*, which abbreviates *q*(**β**, **s**). Furthermore, *η_j_* and *ζ_j_* denote the posterior mean for the effect size and squared effect size, respectively.

Given proposed distribution family *q*, the variational inference scheme proceeds by tuning its parameters to approximate the posterior. The measure of discrepancy between the proposed distribution and the true posterior that we optimize in this work is the Kullback-Leibler (KL-) Divergence [43],

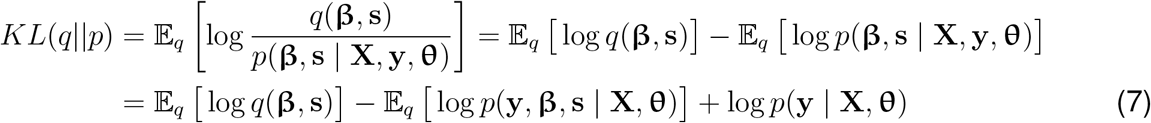

The first term in (7) is the negative entropy of the proposed distribution, whereas the second term is the expectation of the complete log-likelihood with respect to proposed density [43]. The last term is the log marginal likelihood or the model evidence, the normalizing factor that makes the inference intractable. In this setting, the model evidence is a constant with respect to the variational distribution and this fact motivates the use of a surrogate objective, known as the Evidence Lower BOund (ELBO) of the log marginal likelihood [39, 40, 43],

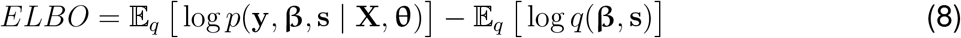

The ELBO combines the expectation of the complete log-likelihood and the entropy of the variational distribution, thus balancing approximation quality against model complexity [39]. Given the model formulation outlined above, the ELBO takes the following form [48, 49],

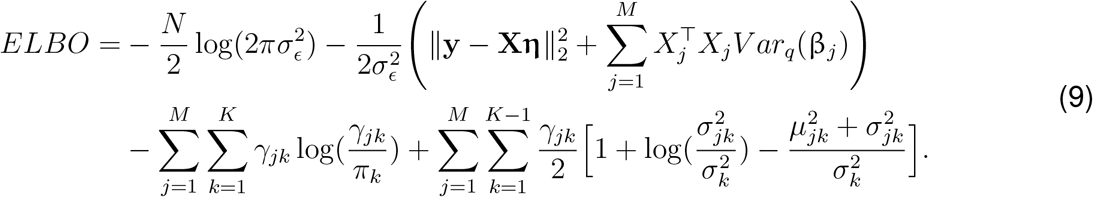

where 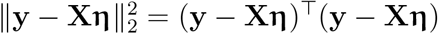. Given the ELBO as the objective function to maximize, we can derive a coordinate ascent Variational EM (VEM) algorithm to learn the variational parameters, as well as the fixed hyperparameter 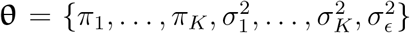 [48, 49, 99]. As detailed below, we update the variational parameters in the E-Step and the hyperparameters in the M-Step, both with the aim of maximizing the ELBO. This process is repeated until convergence.

#### S1.3 Variational EM algorithm

##### The E-Step

In the E-Step, we update the variational parameters for each SNP *j* and mixture component *k* in an iterative fashion, using closed-form updates that can be obtained by taking the partial derivatives of the ELBO with respect to target parameters and solving for the roots [48, 49]:

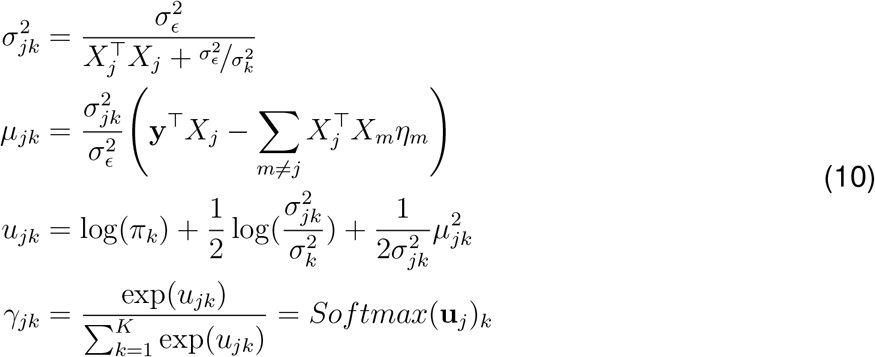

The *Softmax* function applied to the intermediate vector **u**_*j*_ arises from the need to obtain properly normalized posterior densities [39], such that the mixture probabilities for each SNP *j* sum to 1. For the null component (i.e., the spike component), by taking the limit as the prior variance 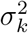 approaches zero 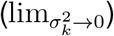 over the update equations above, we see that, as expected, 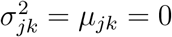 and *u_jk_* = log(*π_k_*).

##### The M-Step

In the M-Step, we update the hyperparameters of the model **θ** to maximize the ELBO as the approximate log marginal likelihood [49]. Similar to the setup in the E-Step, this is done by taking the partial derivatives of the ELBO w.r.t. each hyperparameter *θ_i_* and solving to obtain:

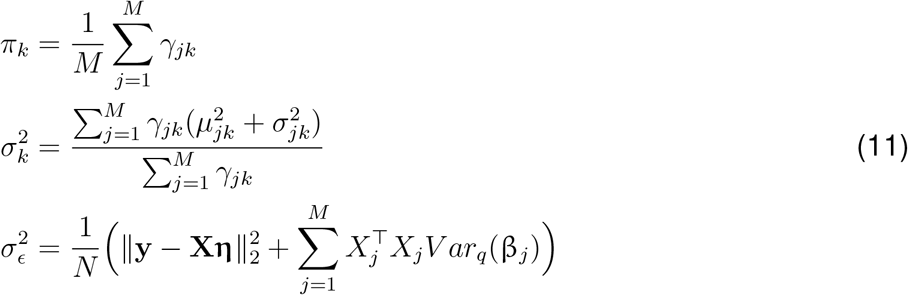

#### S1.4 Expressing the model in terms of GWAS summary statistics

The model formulation above and the resultant closed-form update equations require access to individual-level GWAS data, which can potentially limit its scalability and applicability in many practical settings. An important contribution of this work, therefore, was to re-frame the various components of the model such that it only requires GWAS summary statistics for training [35].

Concretely, by examining the closed-form updates in Eq. (10) and Eq. (11) as well as the ELBO definition in Eq. (8), we see that the individual-level data factors in the updates for the variational parameters 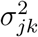, *μ_jk_*, the residual variance 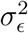, as well as the ELBO. Under the assumption that both the phenotype vector **y** and the genotype matrix **X** have been standardized column-wise to have zero mean and unit variance, the following equivalences hold:

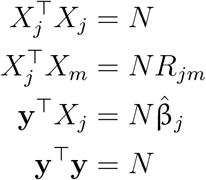

where *N* is the GWAS sample size, **R** is the *M* × *M* Linkage Disequilibrium (LD) matrix that records the pairwise Pearson Correlation coefficient between all pairs of SNPs *j* and *m*, and 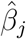 is the standardized marginal GWAS effect size. In this setup, the matrix **R** can be derived from an appropriately-matched reference panel [35]. In many GWAS applications and software implementations (e.g. `plink2` [100]), the genotype matrix is not standardized. To deal with this in practice, we use the pseudo-correlation estimate from Mak et al. (2017) [25] to obtain standardized effect sizes,

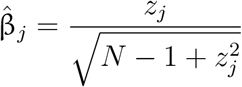

where *z_j_* is the marginal GWAS z-score of SNP *j*. With this in hand, we can write the update equations for the variational parameters in terms of GWAS summary statistics:

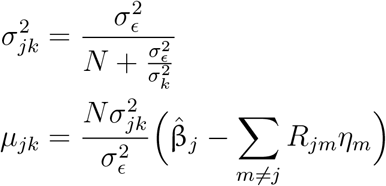

Similarly, in the M-Step, the update for the residual variance, 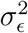, can be written purely in terms of GWAS summary statistics as well:

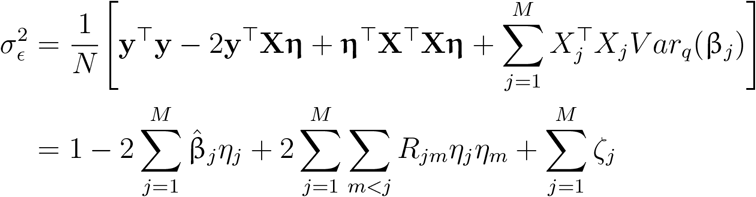

Finally, and using the same expansion for the squared loss as above, we can re-write the objective, the ELBO in Eq. (8), in terms of GWAS summary statistics only,

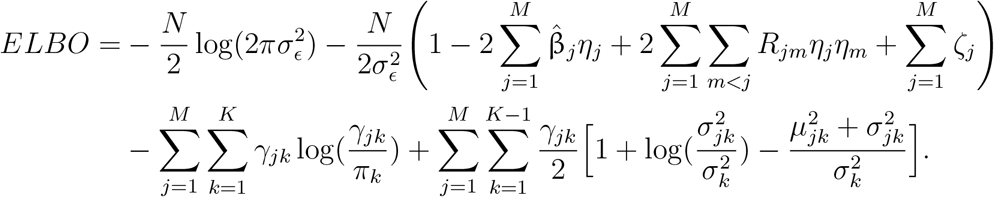

These new update equations form the backbone of the proposed VIPRS model and the new summary statistics-based EM framework is summarized in Algorithm 1.

#### S1.5 Implementation details

##### S1.5.1 Initialization

The Variational EM algorithm for approximate posterior inference is prone to entrapment in local optima, especially in regimes with severe multi-collinearity as is the case with genotype data [49, 72, 104]. Thus, finding an approximate solution that is close to the global optimum requires a well-informed and data-driven parameter initialization strategy. This is particularly important for the hyperparameters of the model, 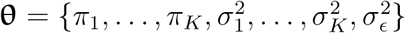, since they are fixed in the generative process outlined above and affect the update equations of all SNPs. For the hyperparameters of the model, we employ the following strategy for initialization:

- For the **mixing proportions** *π_k_*, we draw the overall proportion of causal variants, 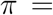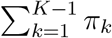, from a uniform between [0.005, 0.1], informed by empirical estimates for a large number of complex traits [10, 11]. Given this random draw, we set the proportion of variants that belong to the null component to *π_K_* = 1 − *π*. The non-null mixture proportions are initialized randomly using a Dirichlet distribution with order *K* − 1 and concentration parameters set to *α_k_* = 1 for all *k* ∈ {1,…, *K* − 1}. Then, the sampled probabilities are multiplied by *π* to ensure that the overall mixing proportions sum to 1.
- For the **residual variance** 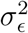, we leverage a fast and simple estimator for the SNP heritability or proportion of variance explained by the SNPs included in the model. For the simple estimator of heritability, we use the formula from [4],

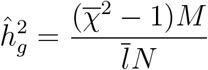

where 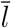 is the average LD score, 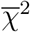 is the average marginal Chi-Squared statistic, and *N* is the GWAS sample size, and *M* is the number of SNPs included in the model. Given this estimate of the heritability, we initialize the residual variance as 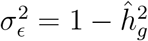.
- For the **prior variances on the effect size** 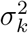, we follow the SBayesR model specification where the different Gaussian components are on different scales, e.g. 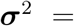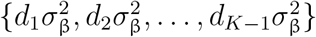 [6]. In our model initialization, we set multiplier to be *d_k_* = 10^−(*k*−1)^, so that the first component has variance 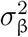, the second component has variance 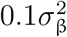, the third component has variance 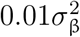, etc. The shared parameter 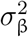 is initialized to be,

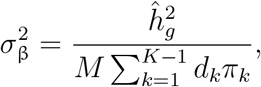

where 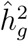 is the naïve SNP heritability estimate, *M* is the number of SNPs included in the model, *d_k_*’s are the multipliers, and *π_k_*’s are the mixing proportions described earlier. In the case of a spike-and-slab prior where *K* = 2, it is easy to see that this simply reduces to the common assumption that 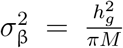, where *π* is the overall proportion of causal variants.

**Algorithm 1:**
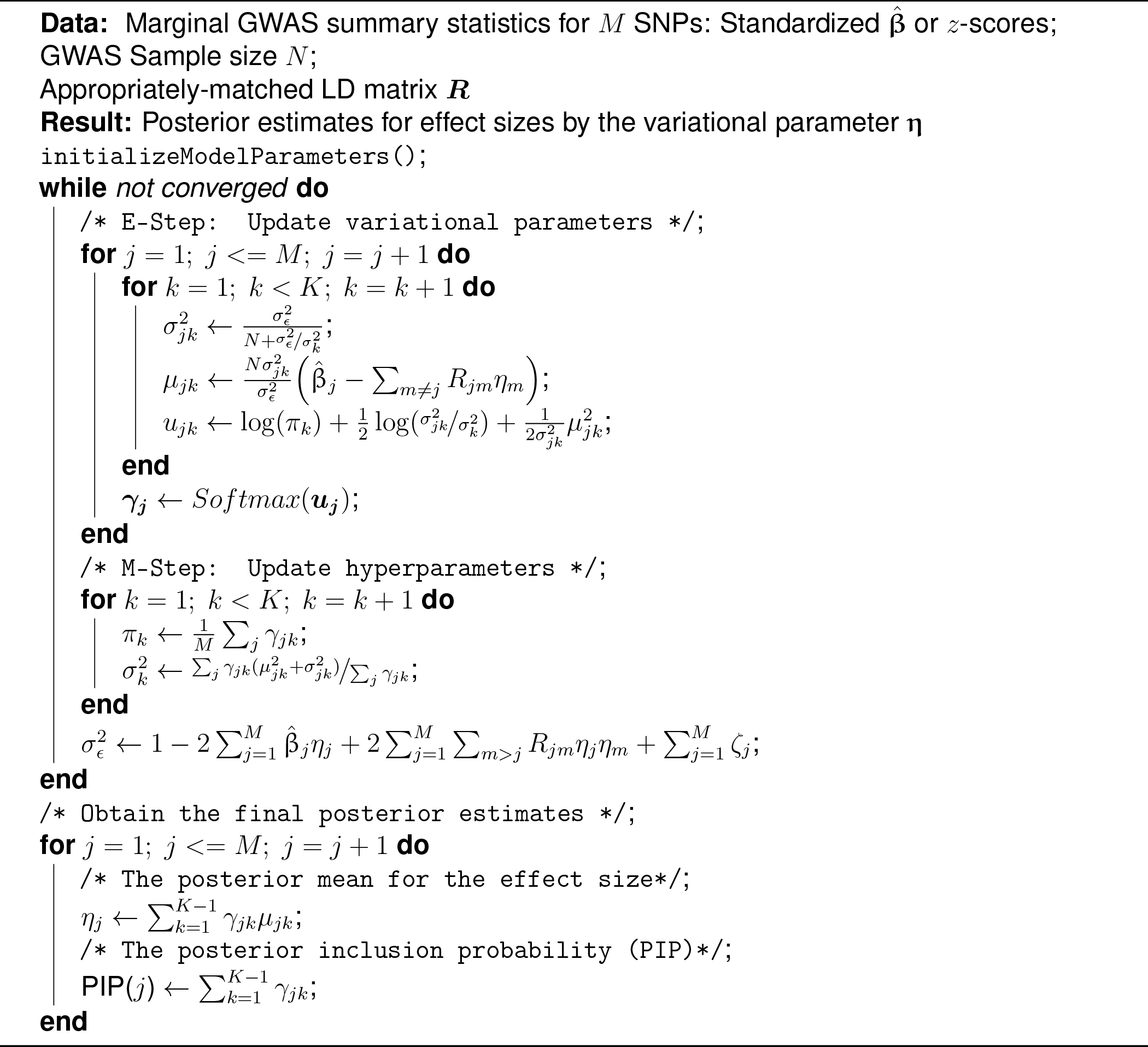
Variational Expectation-Maximization (VEM) algorithm for Bayesian PRS inference with GWAS summary statistics

Once the hyperparameters have been initialized, we initialize the variational parameters *γ_jk_, μ_jk_*, 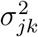 for each SNPs *j* ∈ {1,…, *M*} and each mixture component *k* ∈ {1,…, *K*}. The posterior means for all the mixture components, *μ_jk_*∀*k* ∈ {1,…, *K*}, are simply initialized to be zero. The posterior mixing proportions *γ_jk_*’s, on the other hand, are initialized according to the corresponding priors, i.e. *γ_jk_* = *π_k_* for all *j* and *k*. Finally, the posterior variances 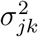 ’s are initialized according to Equation 10, taking the initial values for the prior residual variance 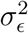 and prior variance 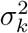 as input.

##### S1.5.2 The *Alpha* prior on the effect size

In studies of the genetic architectures of complex traits, it has been observed that there is a non-linear dependence between the minor allele frequency (MAF) of a genetic variant and its effect size [10, 13, 14]. This MAF-dependent architecture of complex traits is usually modelled with what is known as the “alpha” prior, where the effect size of a SNP is assumed to be drawn from a density where the variance is related to its frequency in the population, e.g.,

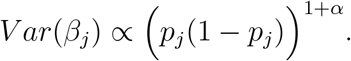

Here, *p_j_* is the minor allele frequency of SNP *j* and *α* is a free parameter that determines the level of dependence between the effect size and the allele frequency. We sought to examine whether incorporating such a prior into the `VIPRS` framework meaningfully improves prediction accuracy. To this end, we derived the `VIPRSAlpha` model, which merges the spike-and-slab prior with the alpha prior, resulting in a mixture density that takes the following form:

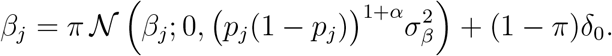

This prior differs from the spike-and-slab in the sense that the per-SNP heritability of causal variants can vary depending on their allele frequency and on the parameter *α*. However, it is more general and in fact reduces to the standard spike-and-slab prior when *α* = −1.

Given a certain *α* value, it is straightforward to show that the update equations in the E-Step (Equation (10)) are not affected by this change, except that we need to replace all instances of 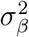 with 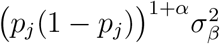 for each SNP *j*. By the same token, this also applies to the closed-form updates for proportion of causal variants *π* and the residual variance 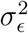 in the M-Step (Equation (11)). However, for the prior variance on the effect size 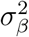, the update equation becomes:

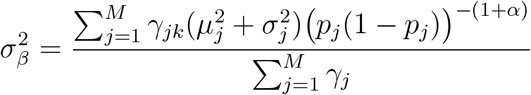

Given this formulation, however, we still need a way to set the unknown parameter *α*. In our software implementation, we allow the user to either set a fixed value for alpha (e.g. *α* = −0.25) or infer its value in the M-Step by optimizing with respect to the ELBO. In the latter case, due to the non-linearity, we use the `L-BFGS-B` optimizer in `scipy` [105] to tune the value of *α*, where the objective function that we minimize is the negative ELBO plus a quadratic penalty term that is equivalent to the standard normal prior imposed on *α* by [10]: 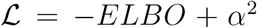. In our experiments, (**Supplementary Fig.** S4 and S5), we optimized the value of *α* in the M-Step.

##### S1.5.3 Hyperparameter tuning strategies

Hyperparameter tuning is important in the Variational EM framework outlined above. Previous work has shown that the EM framework for this model is sensitive to parameter initialization and, therefore, may get stuck in local optima, especially in the presence of strong multicollinearity [49, 72, 104]. This difficulty is compounded by the fact that the model uses point estimates for these hyperparameters, obtained by maximizing the surrogate objective, the ELBO. These maximum likelihood estimates of the hyperparameters do not capture the true uncertainty around them and may lead the model to overfit to the training data [71].

As a remedy, we explored three other strategies for setting or tuning the hyperparameters of the model. These strategies can be deployed in conjunction with the EM framework, or in a hybrid manner. For instance, the mixing proportions *π_k_* can be optimized via cross-validation while the remaining parameters are updated as before in the M-Step [48, 49, 99]. Some of these strategies require access to a held-out validation dataset where we can directly measure the predictive performance associated with each hyperparameter setting.

The three hyperparameter tuning strategies are:

###### Grid Search (GS)

Grid search is a popular strategy for tuning model hyperparameters that has been shown to result in improved predictive performance for a variety of PRS methods [25–27, 30]. This strategy involves two steps that the user can control: **(1)** Grid construction and **(2)** Specifying model selection metric.

For grid construction, the user can choose which hyperparameters to optimize via cross-validation, including the overall proportion of causal variants *π*, the residual variance 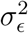, and the prior variance on the effect size 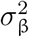. For each hyperparameter, the user specifies the number of values to test as well as the scale of the grid points. For instance, for the proportion of causal variants *π*, we test 30 equidistant points from 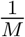 to 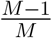 on a log_10_ scale. If the user provides an estimate for any of the hyperparameters, the software can generate a “localized” grid. For example, given a heritability estimate 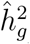, the grid for the residual variance 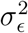 can be multiples of the value provided, e.g. 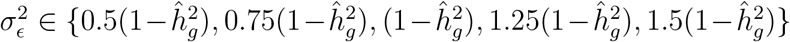 [30]. Once the grid has been specified, the `VIPRS-GS` model is trained, in parallel, for each hyper-parameter combination. The hyperparameters that are not in the grid are updated using the maximum-likelihood estimates derived in Eq. (11). In our experiments, we found that performing grid search over the overall proportion of causal variants *π* with 30 grid points results in a good trade-off between predictive performance and computational efficiency.

In order to select the best hyperparameter combination, we experimented with **three** model selection criteria. The first metric is the prediction *R*^2^ (or AUPRC for case-control phenotypes) on a held-out validation dataset [25, 27, 30]. This approach is generally robust and less prone to over-fitting [61], though it requires that the user has access to an independent validation set with both genotype and phenotype data. It also tends to be computationally intensive for larger validation datasets. As alternative, following earlier work in this area, we also experimented with using pseudo-validation as a criterion for model selection [25, 29, 32]. Given external validation summary statistics, the pseudo-validation criterion, expressed in terms of the Pearson correlation coefficient or coefficient of determination *R*^2^, can be written as:

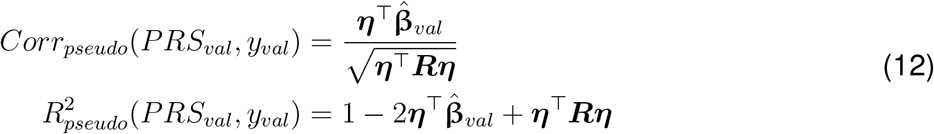

where ***η*** is the vector of posterior means for the effect sizes, 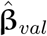 are the standardized marginal effect sizes from the validation set, and ***R*** is the LD matrix. In our software implementation, we simply use the reference LD matrix used during training as proxy for LD in the validation set.

Finally, as a third metric for model selection, we used the training ELBO. The ELBO is a lower bound on the marginal log-likelihood and it balances model fit against model complexity [39, 40]. Given that the standard EM algorithm can get stuck in local optima, exploring many modes of the model using grid search can, in principle, get us closer to the global optimum.

###### Bayesian Model Averaging (BMA)

A different approach for dealing with the unknown hyper-parameters, proposed by [49], is to integrate them out using importance sampling. Concretely, similar to the grid search setup, the idea consists in fitting `VIPRS` with a grid of reasonable values for the hyperparameters. However, instead of selecting the model with the best ELBO or best predictive performance on a validation set, we average the posterior parameter estimates using the importance weights *w*(**θ**^(*g*)^) [49]:

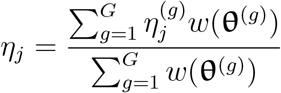

where 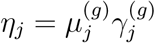 and 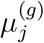 and 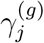 are the expected causal effect and causal indicator, respectively, given the hyperparameters **θ**^(*g*)^. We followed earlier works and take the estimate for the ELBO at convergence as a proxy for the weights *w*(**θ**^(*g*)^)) [48, 49, 99].

###### Bayesian Optimization (BO)

In Bayesian Optimization, we assume that there is an underlying black-box function *f*(**θ**) that takes the hyperparameters as input and outputs a certain score that we wish to optimize, such as the training ELBO or the validation *R*^2^ [62]. This unknown function is modeled with a Gaussian Process (GP) prior, which allows for exploring the hyper-parameter space efficiently while accounting for uncertainty in a principled manner. The other component in this framework is the acquisition function, a heuristic that maps from the GP posterior to information about the most promising regions in hyperparameter space [62, 98]. In our experiments, we used the `scikit-optimize` python package to perform this optimization, with `gp_hedge` as the default acquisition function. By default, the optimizer is allowed to sequentially evaluate up to 20 points in the space of hyperparameters.

In our main experiments, we used Bayesian Optimization to search over the hyperparameter *π*, while updating the remaining remaining hyperparameters using their maximum likelihood estimates, as before [48, 99]. The sequential nature of standard Bayesian optimization procedures makes it slow in practice, compared to e.g. grid search where we can explore different models in parallel. This could potentially be addressed in future work by using parallel implementations of Bayesian optimization [106].

##### S1.5.4 Efficient access and multiplication with the LD matrix

A major bottleneck in the coordinate ascent procedure outlined above is the storage and multiplication with the Linkage-Disequilibrium (LD) matrix ***R*** or its columns. In our software implementation, the matrix ***R***, whether banded, shrunk or dense, is stored on disk in a compressed and chunked array format known as `Zarr` arrays in `python`. During model fitting, if the per-chromosome matrix does not fit in memory, the software is designed to access it chunk by chunk in order to update the model parameters while managing limited memory resources.

While the `Zarr` library provides fast multi-threaded read and write access to the chunked arrays, it still incurs a heavy computational burden compared to arrays that are available in memory. Therefore, in order to get optimal speed and efficiency, ideally we need to minimize read access for the LD matrix when stored on disk. In a naïve implementation of the coordinate ascent procedure outlined above, we would need to access the LD matrix at least 3 times: **(1)** To update the variational parameters *μ_jk_*’s, **(2)** to update the residual variance 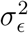, and **(3)** to compute the ELBO, our surrogate objective.

Here we will outline a procedure similar to the “right-hand” updating scheme of SBayesR [6] that keeps track of a per-SNP term, denoted by *q_j_*, which makes it possible to access the LD matrix only once per iteration. We define the *q*–factor for SNP *j* as:

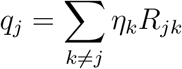

In words, *q_j_* is the sum of the posterior mean for the effect size of all neighboring variants, weighted by their Pearson correlation coefficient with the focal SNP *j*. In our implementation, we initialize all the *q_j_* s to zero and then update the them iteratively during the E-Step. In particular, as we cycle through the SNPs and obtain a new estimate for the posterior mean 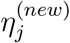, we compute a partial update to the *q* factors for all SNPs in the neighborhood of *j*:

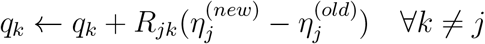

Here, ← denotes the assignment operator, with 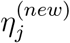 and 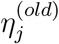 being the new and old posteiror mean estimates for the effect size of SNP *j*, respectively. The benefit of keeping track of this term is that we can then use it when updating the residual variance 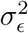 or computing the ELBO without unnecessarily accessing the LD matrix again. The update for residual variance 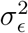 becomes:

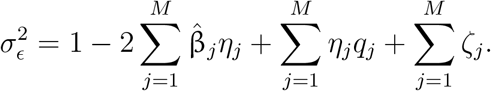

Similarly, the ELBO can be expressed in terms of the q-factor as:

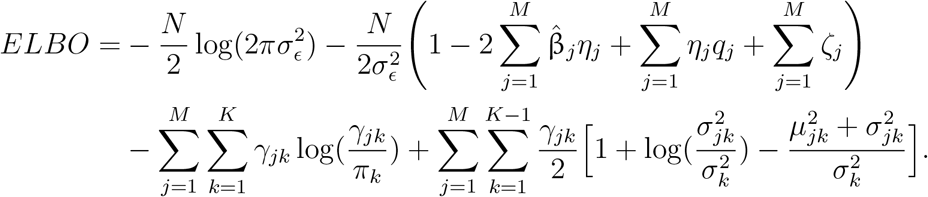

##### S1.5.5 Heritability estimation

An important quantity to help diagnose model fit and quality of posterior approximation is the narrow-sense heritability, or proportion of additive variance explained (PVE) by the SNPs included in the model [107]. In the context of the linear model outlined in Equation 2, SNP heritability is defined as:

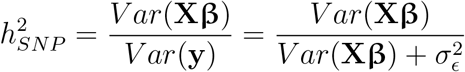

where *V ar*(**Xβ**) denotes the genotypic variance and *V ar*(**y**) is the phenotypic variance, which we assume to be a linear sum of the genotypic and residual variances. Therefore, given this formulation, estimating SNP-heritability is equivalent to the task of estimating the genotypic and residual variances under the posterior distribution [6, 48].

In our model, the residual variance 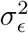 is a point estimate that we update in the M-Step of the Variational EM algorithm and that point estimate is used in the denominator above to compute an estimate of SNP-heritability. As for the genotypic variance, assuming that both **X** and **β** are random, we can write:

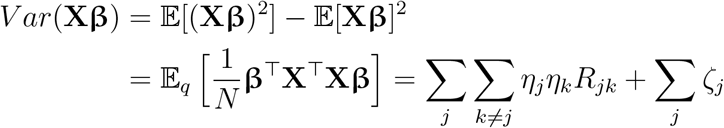

where the expectation is taken with respect to all the random components of the model. The second equality follows from our assumption that the genotype matrix is standardized to have mean zero and unit variance. The above estimator for the genotypic variance under the variational posterior can be simplified and expressed in terms of the *q*–factor described previously,

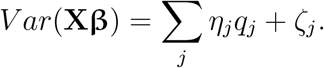

Previous work has noted that the Variational approximation to the posterior tends to systematically underestimate SNP heritability or PVE [48, 49]. In our experiments, we saw that the estimated heritability is accurate for sparse genetic architectures, though it tends to be systematically underestimated for highly polygenic traits (**Supplementary Fig.** S8).

#### S1.6 A heuristic test of mismatch between GWAS summary statistics and LD reference panel

Summary statistics-based PRS methods can be sensitive to heterogeneities between GWAS summary statistics and the LD reference panel [8, 30, 76]. For some Bayesian methods, this mismatch can result in unpredictable behavior, with the posterior mean for the effect sizes exploding in magnitude and the estimated SNP-heritability exceeding 1 in some circumstances^1^. We observed that these inference difficulties can arise in the context of the `VIPRS` model, with the model objective, the ELBO, rapidly diverging after a few iterations. Thus, we sought ways to detect and eliminate strong inconsistencies between the GWAS cohort and the LD reference panel.

In general, if the requisite data is available, we encourage users to run principled and comprehensive tests of heterogeneity, such as the `DENTIST` method of Chen et al. (2021) [77]. However, in case this is not feasible, due to e.g. using precomputed LD matrices without direct access to the individual-level data of the reference panel, we designed a fast, heuristic and noisy estimator of the `DENTIST` p-value (***P***_DENTIST_) that leverages only pre-computed summary statistics. As explained below, our experiments indicate that this heuristic approach can help inference robustness by identifying problematic SNPs.

The heuristic test of mismatch works as follows: For every variant *i* included in the analysis, we randomly sample a fixed number of neighbors *G* with replacement. The neighbors in this case are defined by the pre-computed LD matrix. For instance, for the windowed estimator of LD, the neighbors are only SNPs that are within a pre-specified window around the focal SNP. Then for every sampled neighbor *j*, we compute the *T_d_* statistic as [77],

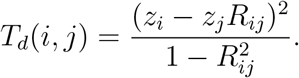

The final *T_d_* statistic for variant *i*, then, is the average across the SNP’s sampled neighbors 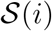,

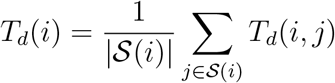

Given these noisy estimates of the *T_d_* statistic, we then compute a p-value, assuming that it remains roughly Chi-squared distributed with 1 degree-of-freedom [77]. If the p-value is less than a Bonferroni threshold of 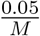, where *M* is the number of variants included in the model, then those variants are filtered out. In practice, we found that a single round of filtering sometimes does not eliminate all inconsistencies. Thus, in our software pipeline, we follow the authors of the `DENTIST` method and apply the filters iteratively until model convergence is achieved or we reach a predefined maximum number of iterations.

#### S1.7 VIPRS contributions and related work

The statistical model underlying the `VIPRS` framework was originally inspired by the work of Titsias and Lázaro-Gredilla (2011) [59] and the notation and formalism employed here are partly based on their model description as well as an important early paper by Carbonetto and Stephens (2012) [49] on using variational inference for Bayesian variable selection in the context of genetics data. Both of these works employed a sparse mixture prior (spike-and-slab) to model high-dimensional data. The latter work, implemented in the software package varbvs [99], was concerned with the problem of fine or association mapping in statistical genetics and demonstrated the scalability of variational inference by applying it to a sample of roughly 5000 individuals and half a million predictors (SNPs) from the Wellcome Trust Case Control Consortium (WTCCC) 1 study [108]. Carbonetto and Stephens (2012) also demonstrated some of the limitations of variational approximations in the context of fine-mapping, especially in the presence of strong multicollinearity or Linkage Disequilibrium (LD) between SNPs. In addition to anticipating and inspiring multitudes of theoretical studies examining the properties of these sparse variational approximations in the statistics literature [72, 109, 110], the varbvs model also inspired a number of extensions in the context of statistical genetics.

Many of the model extensions of `varbvs` were concerned with adding extra layers of interpretability by incorporating genomic annotations and gene pathway information into the sparse prior [51]. This enabled principled enrichment analyses and gene prioritization pipelines. Like the original `varbvs` model, this extension operated on individual-level data, and thus it required access to genotype and phenotype data from the same sample. More recently, with the development of the Regression with Summary Statistics (RSS) likelihood [111], a summary statistics version of this enrichment and prioritization model was developed [52]. Even though the `VIPRS` model did not employ the RSS likelihood, the closed-form parameter updates that we obtain for the spike-and-slab prior are very similar to those found in the `RSS-E` model [52]. These efforts culminated in the recent development of the Biologically Annotated Neural Network (`BANN`) framework, which models the linear and non-linear contributions of genotype to the phenotype in an integrated Bayesian neural network model [48]. In addition to the non-linearities, the `BANN` model also extended the spike-and-slab prior into a sparse Gaussian mixture prior with arbitrary *K* components, which we also tested with the `VIPRSMix` model formulation (this prior is also used in the `BayesR` family of genomic prediction models [6, 24]). The `BANN` model was derived mainly for use with individual-level data, though a summary statistics version was also proposed by the authors. To our knowledge, none of these models have been tested or evaluated in the context of prediction or polygenic risk estimation.

In addition to this line of work, the Bayesian sparse priors of the form employed by `VIPRS` have been used extensively in statistical genetics applications. For instance, Zhou et al. (2013) proposed a model that combines the spike-and-slab prior (BVSR in their terminology) with the Linear Mixed Model (LMM) formulation for the purposes of genomic prediction and heritability estimation [112]. However, the resulting `BSLMM` model operated on individual-level data and used MCMC for posterior inference, thus limiting its scalability in practice. A recent extension of this model (`DBSLMM`), which operates on GWAS summary statistics and uses a Clumping and Thresholding heuristic procedure to deterministically partition variants into small and large effect SNPs, has been shown to be competitive for the task of polygenic prediction [29, 37, 38]. However, unlike `VIPRS` and other Bayesian methods tested here, this model is not sparse, instead assuming an infinitesimal genetic architecture for complex traits.

To our knowledge, the first Bayesian summary statistics-based PRS method that employed the spike-and-slab prior is the `LDPred` model [4], which, with recent software and statistical improvements, remains one of the most competitive PRS methods in the literature [30, 37, 38]. The `LDPred` and `LDPred2` models still employ MCMC for posterior inference, however. Incidentally, in the discussion of the original `LDPred` paper, the authors allude to the work of Carbonetto and Stephens (2012) [49] among others, stating that variational approximations did not seem to perform satisfactorily in their initial experiments [4]. In this context, one of the main contributions of our work was showing that summary statistics-based variational approximations can be highly accurate and competitive with MCMC-based methods.

As alluded to in the main text, `VIPRS` is not the first model to use variational inference for polygenic prediction either from GWAS summary statistics or individual-level data. In addition to the fine-mapping and enrichment analysis pipelines discussed previously, variational approximations have recently been explored by a number of authors for PRS estimation. For example, Zeng and Zhou (2017) employed the mean-field variational inference framework to fit a Dirichlet Process Regression (`DPR`) model to individual-level data [53]. In their work, they showed that variational inference for their model performs less well than MCMC, though it’s generally very competitive with existing baselines. Recently, the `DPR` model was extended to use summary statistics (`SDPR`), though, to our knowledge, the authors only provide MCMC solutions to approximate the posterior for the model parameters [31].

More recently, Zhang et al. (2021) proposed `MegaPRS`, a whole suite of PRS methods that work with individual-level data or summary statistics and can accommodate a variety of heritability models and prior families [32]. For inference, the authors employed a variational algorithm inspired by the BOLT-LMM inference procedure [46] (Doug Speed, personal communications). `MegaPRS` differs from `VIPRS` in the sense that it does not employ a Variational EM framework. In their method, some of the hyperparameters of the model are inferred beforehand using external software (e.g. `SumHer`), some are fixed to reasonable values (e.g. setting the residual variance to 1.), and some are fit to the data using cross-validation (e.g. mixing proportions). To stabilize the variational algorithm, the `MegaPRS` method further restricts the optimization procedure to overlapping windows of 1/8th of a centiMorgan. On the other hand, `VIPRS` implements the full coordinate ascent procedure on a chromosome-by-chromosome basis, cycling through all the SNPs and iteratively updating their variational parameters until convergence. As we showed in our analyses, these modeling choices result in better variational approximations in most scenarios, as attested by the fact the `VIPRS` out-performed `MegaPRS` in most of our experiments.

Finally, concurrent with our work, Spence et al. (2022) proposed the `Vilma` model, a variational inference framework for inferring effect size distributions across ancestrally diverse populations [55]. Unlike `VIPRS`, the `Vilma` model is based on the `RSS` likelihood, thus it can be considered a multi-population extension of the `RSS` and `RSS-E` models, with an arbitrarily flexible sparse Gaussian mixture prior. Another point of distinction between `Vilma` and our method is that whereas `VIPRS` uses a fast coordinate ascent scheme for optimizing the parameters of the proposed variational density, the authors of `Vilma` employ a gradient ascent procedure which necessitates computing the pseudo-inverse of the LD matrix [55]. Thus, `VIPRS` is much faster and more scalable in practice.

### S2 Supplementary Figures

#### S2.1 Predictive performance of summary statistics-based PRS methods on simulated binary traits

**Figure S1:**
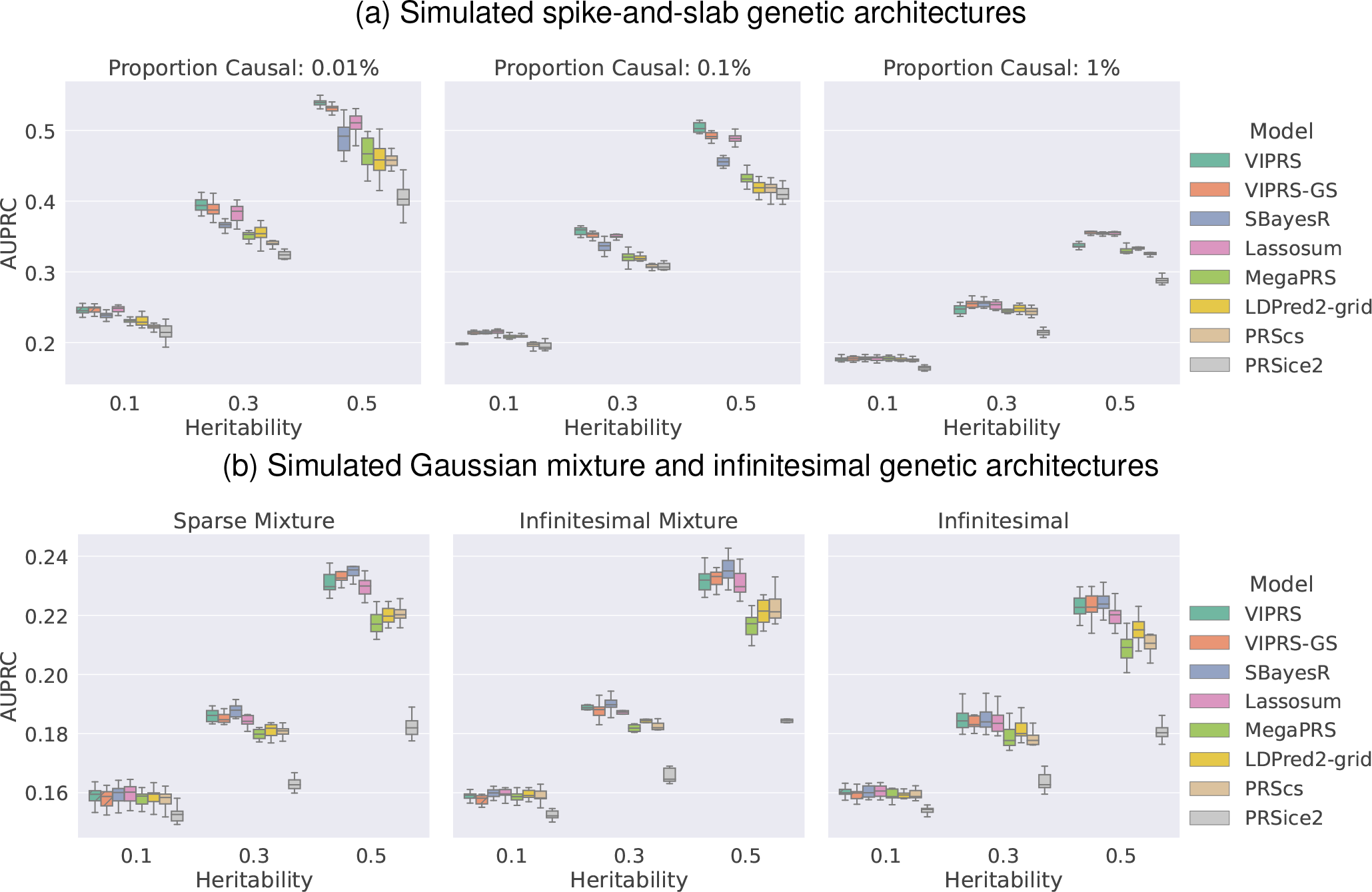
Predictive performance of summary statistics-based PRS methods on simulated binary traits (prevalence 15%) following **(a)** spike-and-slab and **(b)** Gaussian mixture or infinitesimal genetic architectures. The phenotypes were simulated using real genotype data from the White British cohort in the UK Biobank (*N* = 337, 225), leveraging a subset of 1.1 million HapMap3 variants. The simulation scenarios encompass a total of 18 configurations, spanning 6 genetic architectures and 3 values for SNP heritability. For each configuration, we simulated 10 independent phenotypes. Each panel shows results for phenotypes simulated with the pre-specified genetic architecture and each column within a panel shows performance metrics for phenotypes simulated with a pre-specified SNP heritability. The performance metric shown is the area under the Precision Recall curve (AUPRC). The boxplot for each method and simulation configuration shows the quartiles of the AUPRC scores for the 10 simulated phenotypes. The PRS methods shown are our proposed `VIPRS` and `VIPRS-GS` (using grid search to tune model hyperparameters) as well as 5 other baseline models: `SBayesR`, `Lassosum`, `MegaPRS`, LDPred2 (grid), `PRScs`, and `PRSice2` (C+T).

#### S2.2 Performance of VIPRS with different LD estimators and sample sizes for the LD reference panel

**Figure S2:**
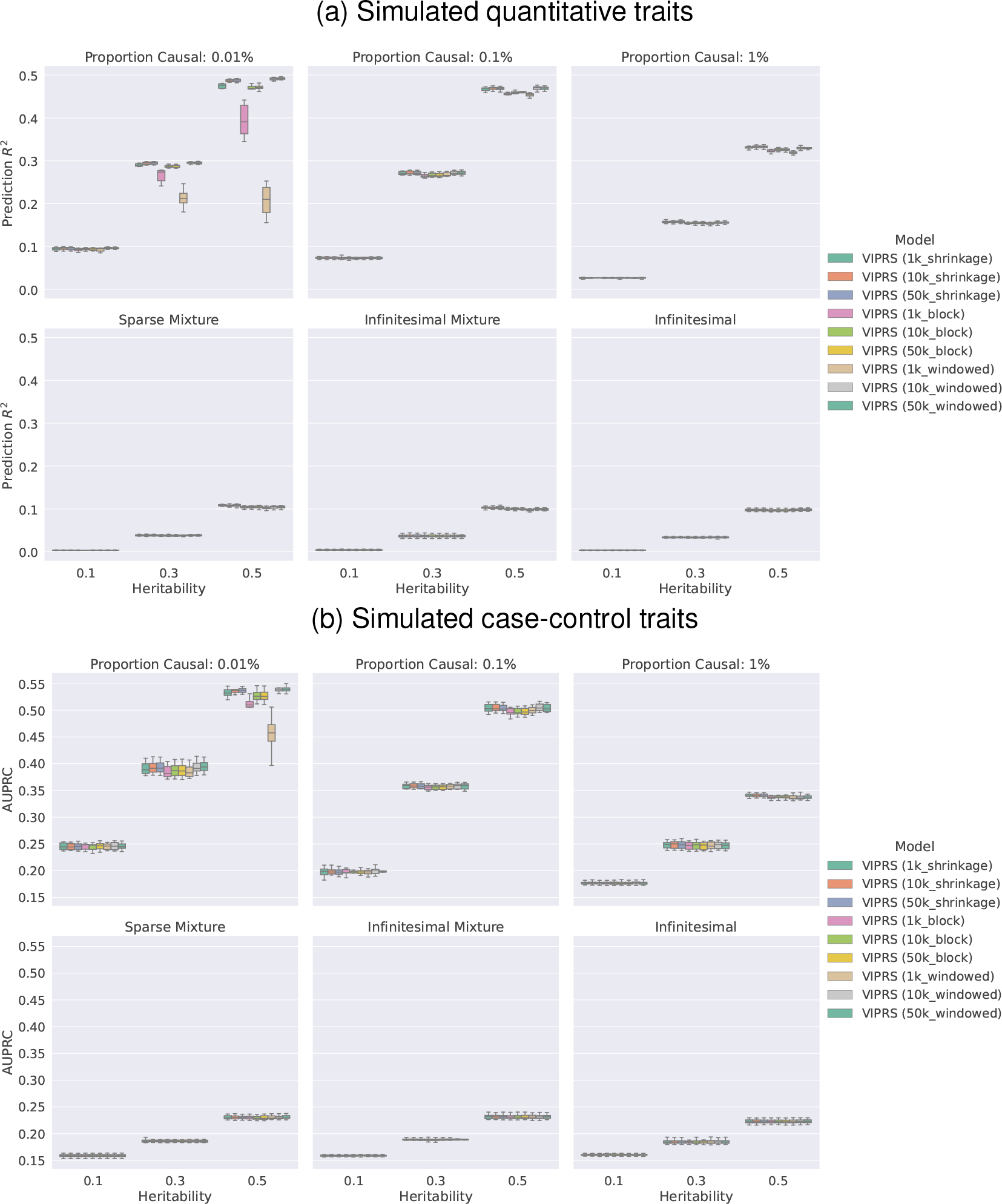
Predictive performance of the `VIPRS` model on simulated phenotypes using different estimators and sample sizes for the LD reference panel. The simulated phenotypes encompass a total of 6 genetic architectures and 3 values for SNP heritability (see **Methods**). Each panel shows results for phenotypes simulated with the pre-specified architecture and each column within a panel shows performance metrics for phenotypes simulated with a pre-specified SNP heritability. The performance metrics are **(a)** incremental prediction *R*^2^ for quantitative traits and **(b)** area under the Precision Recall curve (AUPRC) for binary traits. The boxplot for each method and simulation configuration shows the quartiles of predictive scores for the 10 simulated phenotypes. The parenthesis in each method name indicate the sample size of the LD reference panel (1000, 10000, 50000) as well as the LD estimator (shrinkage, block, windowed).

**Figure S3:**
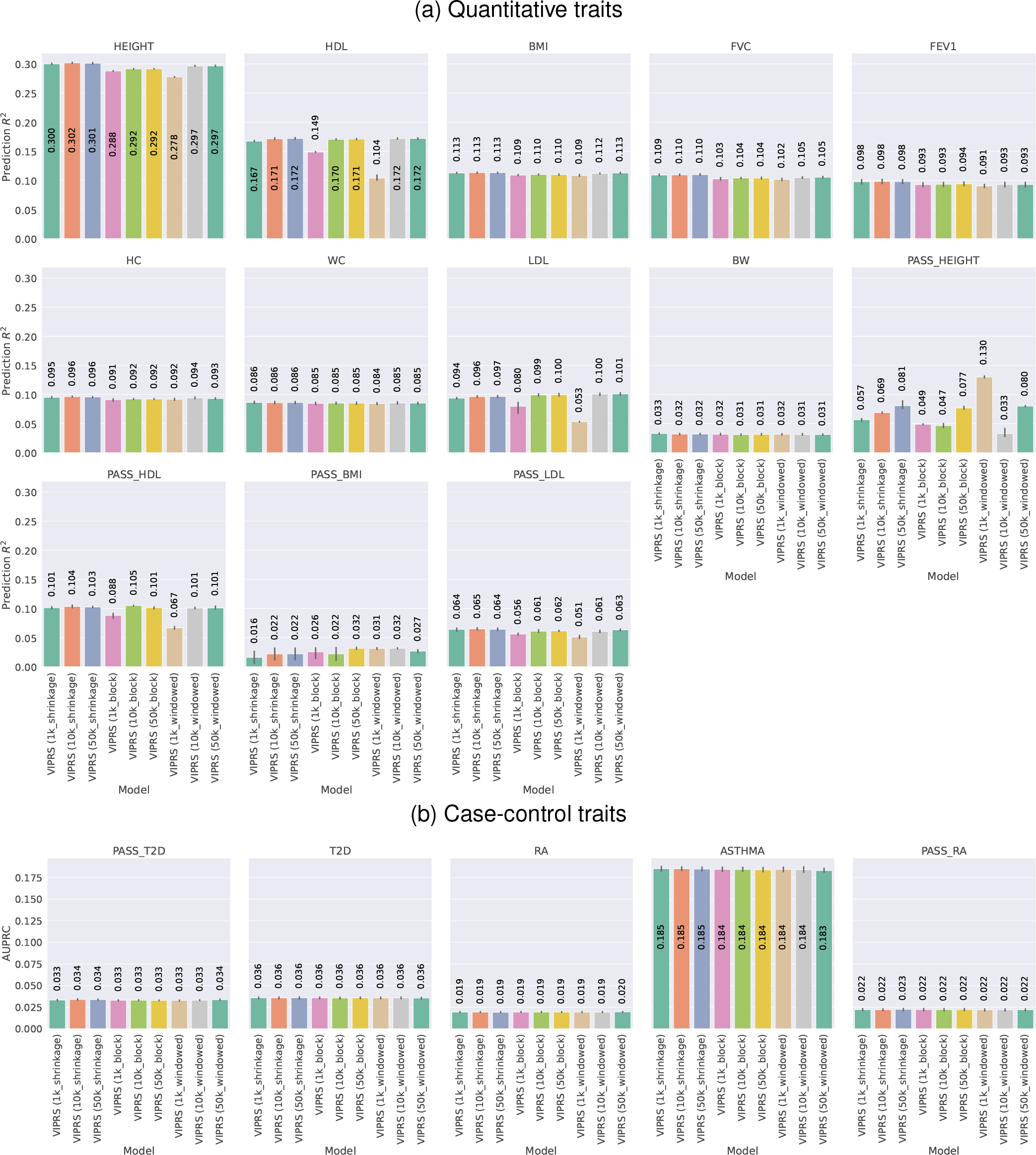
Predictive performance of the `VIPRS` model on real **(a)** quantitative and **(b)** binary phenotypes in the UK Biobank using different estimators and sample sizes for the LD reference panel. Each panel shows the predictive performance, in terms of **(a)** incremental *R*^2^ and **(b)** area under the Precision Recall curve (AUPRC), of each PRS model when applied to a given phenotype. The bars show the mean of the prediction metrics across the 5 folds and the black lines show the corresponding standard errors. The phenotypes analyzed are described in detail in Table 1.The parenthesis in each method name indicate the sample size of the LD reference panel (1000, 10000, 50000) as well as the LD estimator (shrinkage, block, windowed).

#### S2.3 Performance of the VIPRS model with different families of priors on the effect size

**Figure S4:**
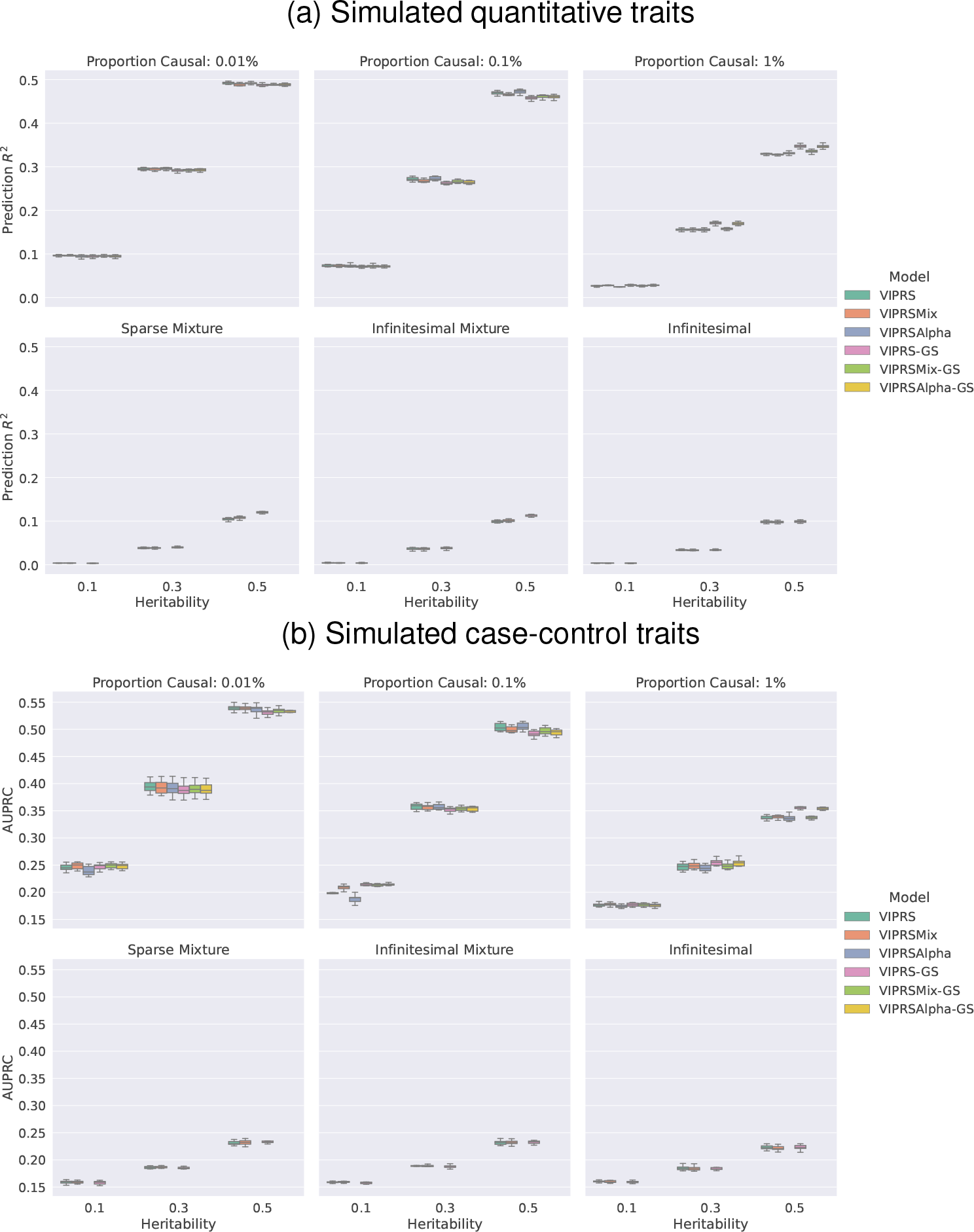
Predictive performance of the `VIPRS` model on simulated phenotypes using different families of priors on the effect size. The simulated phenotypes encompass a total of 6 genetic architectures and 3 values for SNP heritability (see **Methods**). Each panel shows results for phenotypes simulated with the pre-specified architecture and each column within a panel shows performance metrics for phenotypes simulated with a pre-specified SNP heritability. The performance metrics are **(a)** incremental prediction *R*^2^ for quantitative traits and **(b)** area under the Precision Recall curve (AUPRC) for binary traits. The boxplot for each method and simulation configuration shows the quartiles of predictive scores for the 10 simulated phenotypes. The methods tested are `VIPRS` (spike-and-slab prior), `VIPRSMix` (sparse Gaussian mixture prior with four components), and `VIPRSAlpha` (spike-and-slab + Alpha prior). In addition, we show the performance of the grid-search version of the model with the aforementioned three priors (`VIPRS-GS`, `VIPRSMix-GS`, `VIPRSAlpha-GS`), where the search is over the proportion of causal variants *π*.

**Figure S5:**
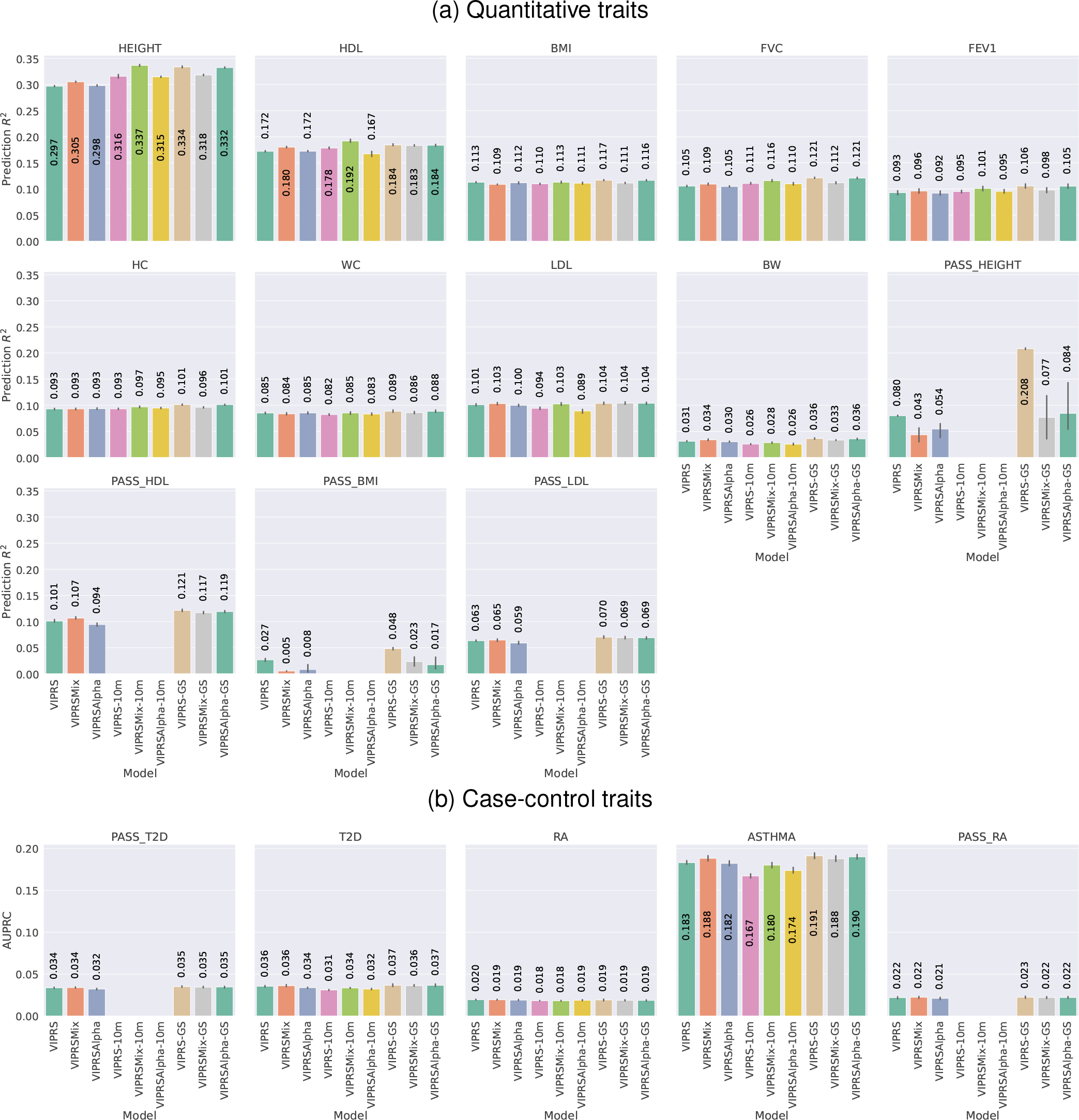
Predictive performance of the `VIPRS` model on real **(a)** quantitative and **(b)** binary phenotypes in the UK Biobank using different families of priors on the effect size. Each panel shows the predictive performance, in terms of **(a)** incremental *R*^2^ and **(b)** area under the Precision Recall curve (AUPRC), of each PRS model when applied to a given phenotype. The bars show the mean of the prediction metrics across the 5 folds and the black lines show the corresponding standard errors. The phenotypes analyzed are described in detail in Table 1.The methods tested are `VIPRS` (spike-and-slab prior), `VIPRSMix` (sparse Gaussian mixture prior with four components), and `VIPRSAlpha` (spike-and-slab + Alpha prior). These methods were evaluate with the HapMap3 SNPs as well as the 9.6 million expanded set of variants. For the latter case, we added the `10m` suffix to the method name. In addition, we show the performance of the grid-search version of the model with the aforementioned priors (`VIPRS-GS`, `VIPRSMix-GS`, `VIPRSAlpha-GS`), where the search is over the proportion of causal variants *π*. **NOTE**: We did not run the methods with 9.6 million variants on external summary statistics since they only contained marginal test statistics for a limited set of SNPs.

#### S2.4 Performance of the VIPRS model with different hyperparameter tuning strategies

**Figure S6:**
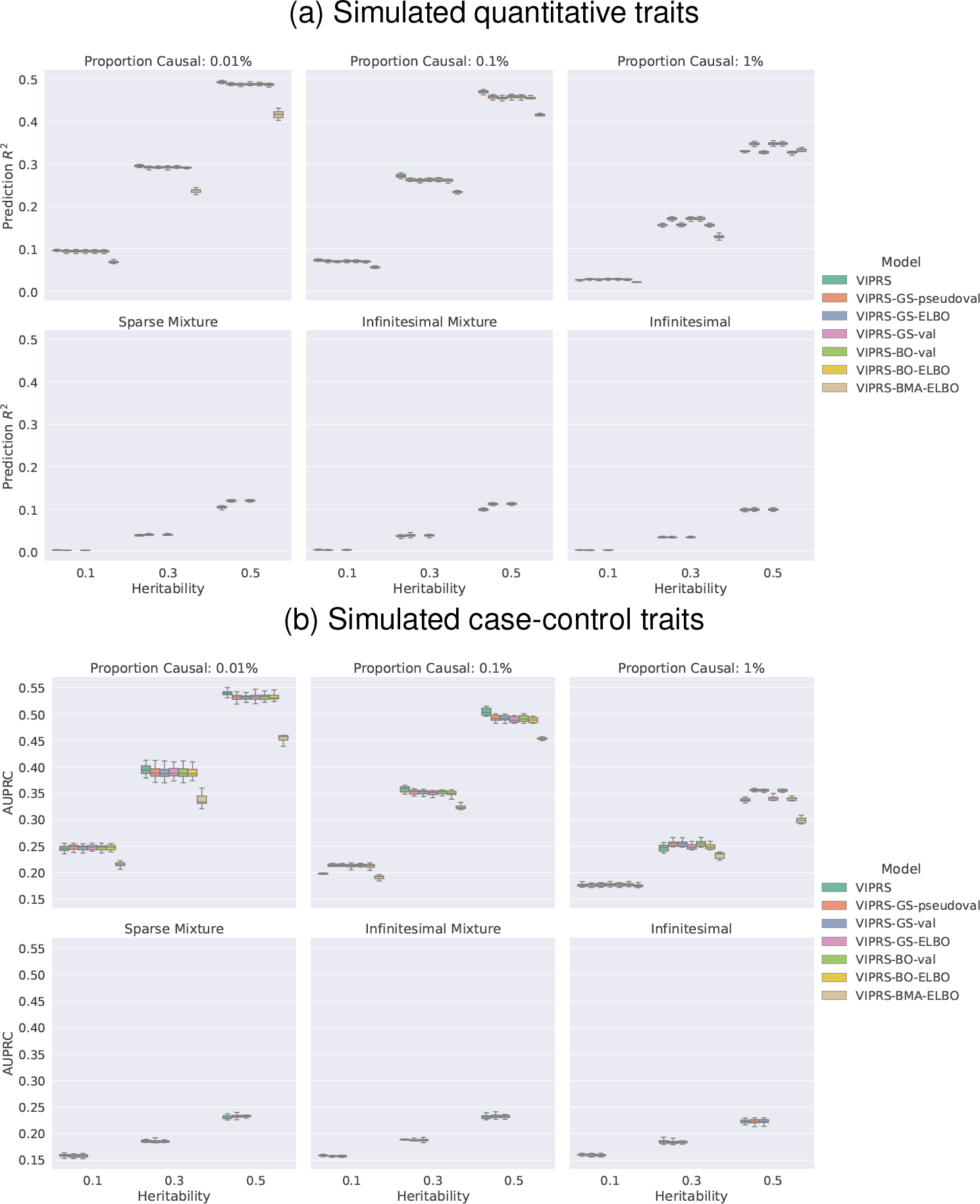
Predictive performance of the `VIPRS` model on simulated phenotypes using different hyperparameter tuning strategies. The simulated phenotypes encompass a total of 6 genetic architectures and 3 values for SNP heritability (see **Methods**). Each panel shows results for phenotypes simulated with the pre-specified architecture and each column within a panel shows performance metrics for phenotypes simulated with a pre-specified SNP heritability. The performance metrics are **(a)** incremental prediction *R*^2^ for quantitative traits and **(b)** area under the Precision Recall curve (AUPRC) for binary traits. The boxplot for each method and simulation configuration shows the quartiles of predictive scores for the 10 simulated phenotypes. The hyperparameter tuning strategies tested are `VIPRS` (Variational EM), `VIPRS-GS-ELBO` (grid search with the ELBO as criterion), `VIPRS-GS-pseudoval` (grid search with pseudo-validation), `VIPRS-GS-val` (grid search with validation), `VIPRS-BO-ELBO` (Bayesian optimization with the ELBO as criterion), `VIPRS-BO-val` (Bayesian optimization with validation), and `VIPRS-BMA` (Bayesian Model Averaging). In these experiments, the hyperparameter search is over the proportion of causal variants *π*.

**Figure S7:**
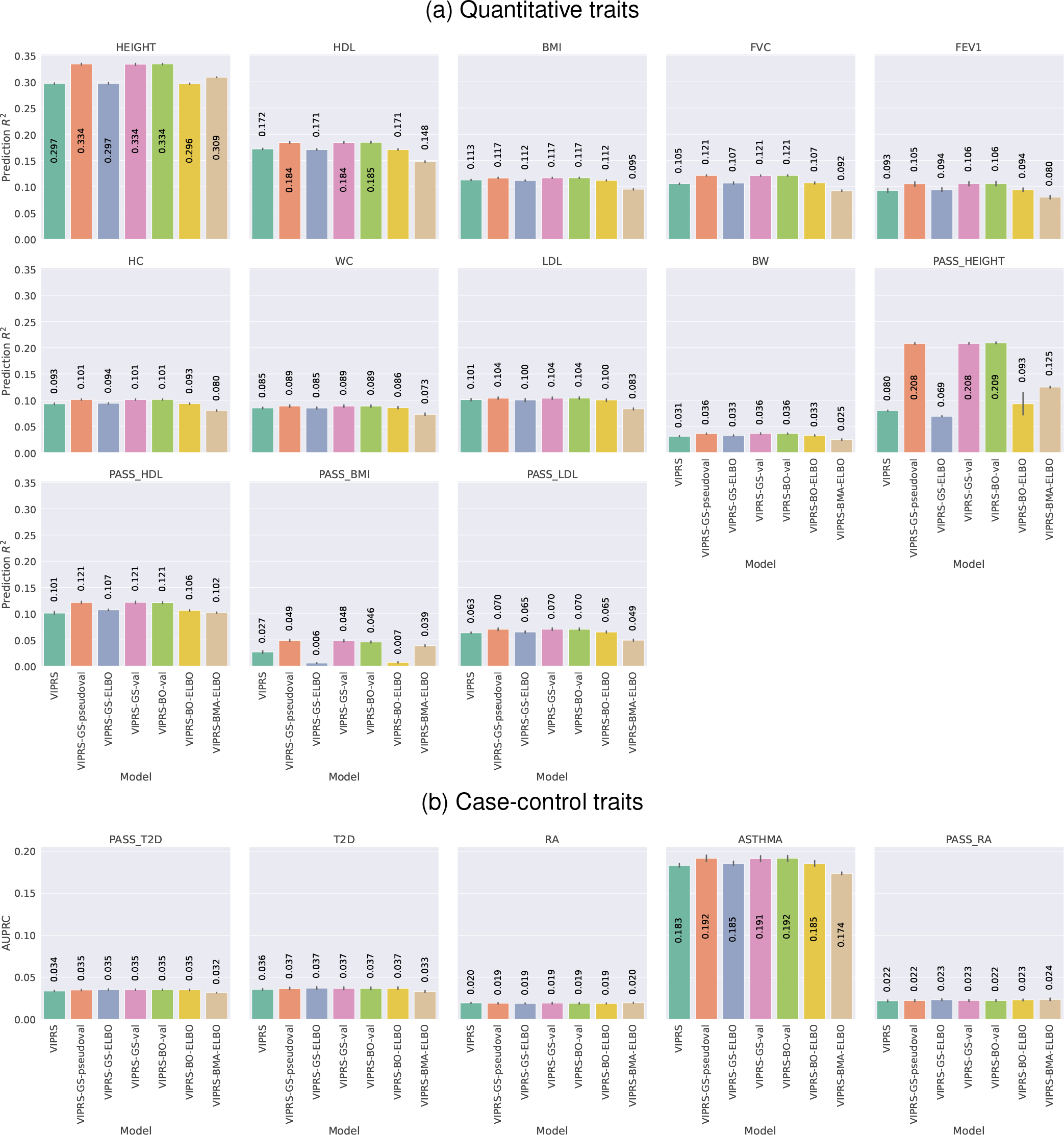
Predictive performance of the `VIPRS` model on real **(a)** quantitative and **(b)** binary phenotypes in the UK Biobank using different hyperparameter tuning strategies. Each panel shows the predictive performance, in terms of **(a)** incremental *R*^2^ and **(b)** area under the Precision Recall curve (AUPRC), of each PRS model when applied to a given phenotype. The bars show the mean of the prediction metrics across the 5 folds and the black lines show the corresponding standard errors. The phenotypes analyzed are described in detail in Table 1. The hyperparameter tuning strategies tested are `VIPRS` (Variational EM), `VIPRS-GS-ELBO` (grid search with the ELBO as criterion), `VIPRS-GS-pseudoval` (grid search with pseudo-validation), `VIPRS-GS-val` (grid search with validation), `VIPRS-BO-ELBO` (Bayesian optimization with the ELBO as criterion), `VIPRS-BO-val` (Bayesian optimization with validation), and `VIPRS-BMA` (Bayesian Model Averaging). In these experiments, the hyperparameter search is over the proportion of causal variants *π*.

#### S2.5 Hyperparameter estimation: SNP heritability and polygenicity

**Figure S8:**
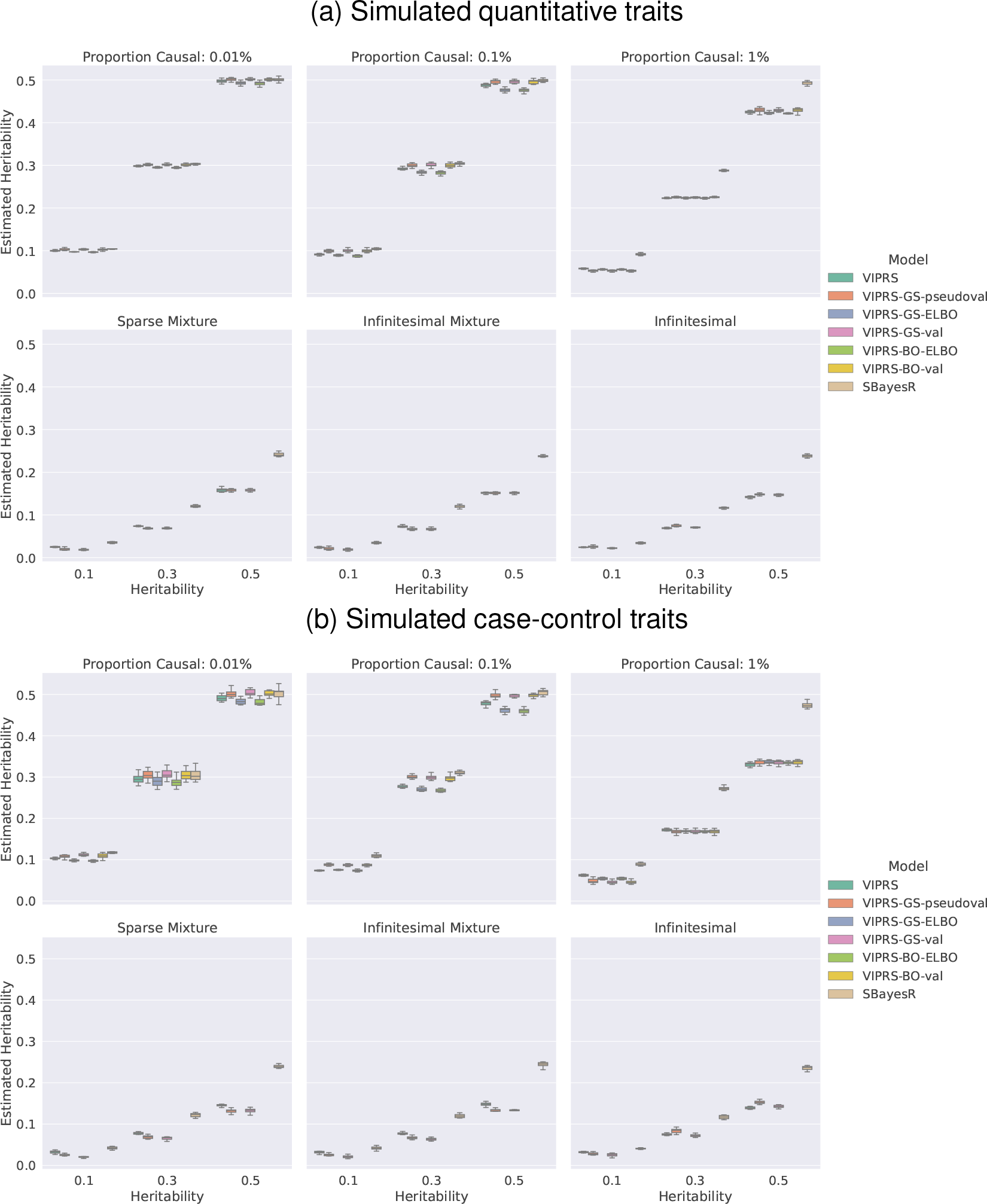
Estimates of SNP heritability or proportion of variance explained (PVE) by summary statistics-based PRS methods for simulated **(a)** quantitative and **(b)** case-control traits. The simulated phenotypes encompass a total of 6 genetic architectures and 3 values for SNP heritability (see **Methods**). Each panel shows results for phenotypes simulated with the pre-specified architecture and each column within a panel shows performance metrics for phenotypes simulated with a pre-specified SNP heritability. The performance metrics are **(a)** incremental prediction *R*^2^ for quantitative traits and **(b)** area under the Precision Recall curve (AUPRC) for binary traits. The boxplot for each method and simulation configuration shows the quartiles of predictive scores for the 10 simulated phenotypes. The methods tested are `VIPRS` (Variational EM), `VIPRS-GS-pseudoval` (grid search with pseudoval-idation), `VIPRS-GS-val` (grid search with validation), `VIPRS-BO-ELBO` (Bayesian optimization with the ELBO as criterion), `VIPRS-BO-val` (Bayesian optimization with validation), and `SBayesR`.

**Figure S9:**
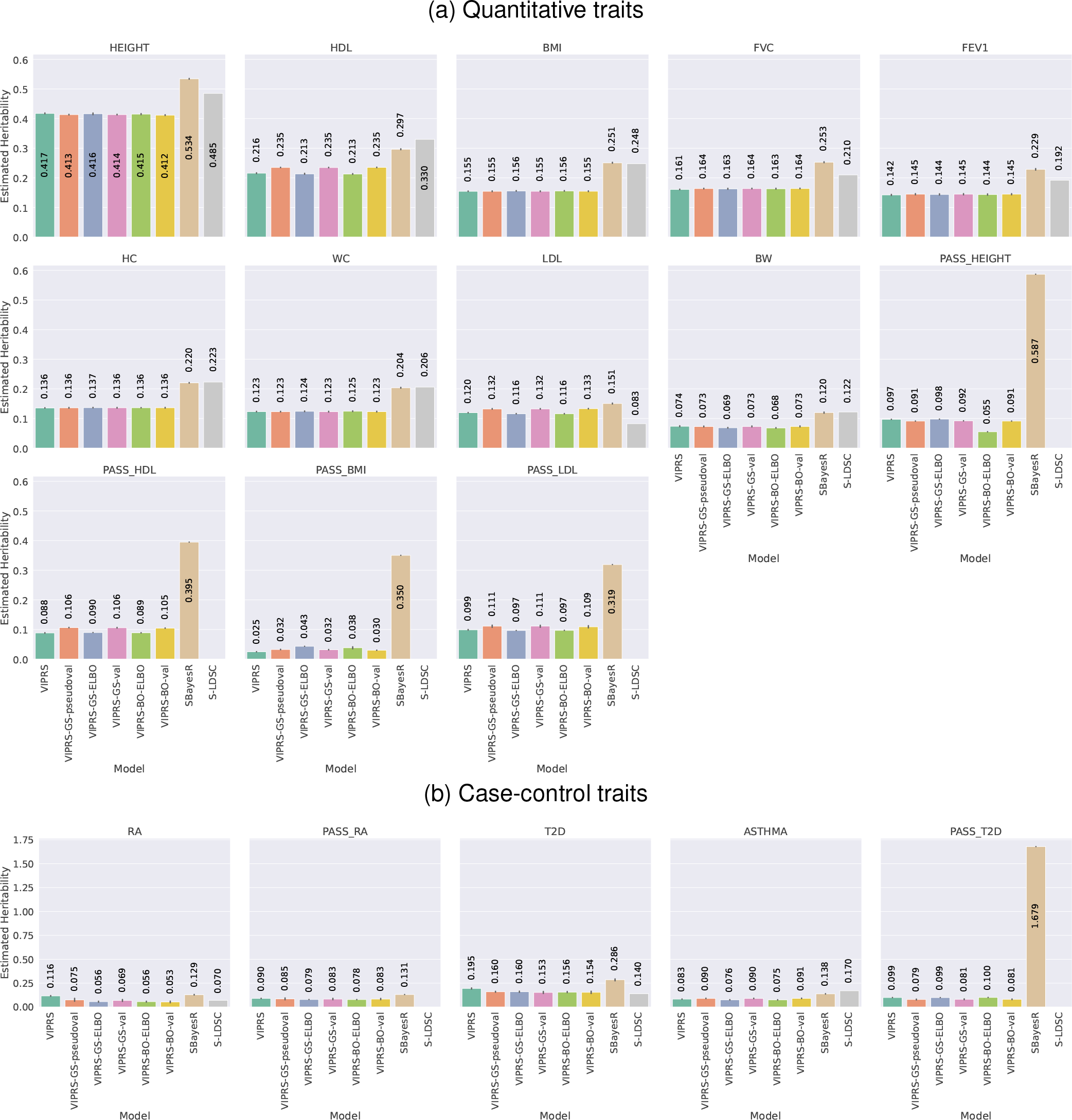
Estimates of SNP heritability or proportion of variance explained (PVE) by summary statistics-based PRS methods for real **(a)** quantitative and **(b)** case-control traits in the UK Biobank. The bars show the mean estimate of SNP heritability across the 5 folds and the black lines show the corresponding standard errors. The phenotypes analyzed are described in detail in Table 1. The methods tested are `VIPRS` (Variational EM), `VIPRS-GS-pseudoval` (grid search with pseudovalidation), `VIPRS-GS-val` (grid search with validation), `VIPRS-BO-ELBO` (Bayesian optimization with the ELBO as criterion), `VIPRS-BO-val` (Bayesian optimization with validation), and `SBayesR`, and, for reference, `S-LDSC` (estimates are taken from Ben Neale lab’s Heritability Browser: https://nealelab.github.io/UKBB_ldsc/h2_browser.html).

**Figure S10:**
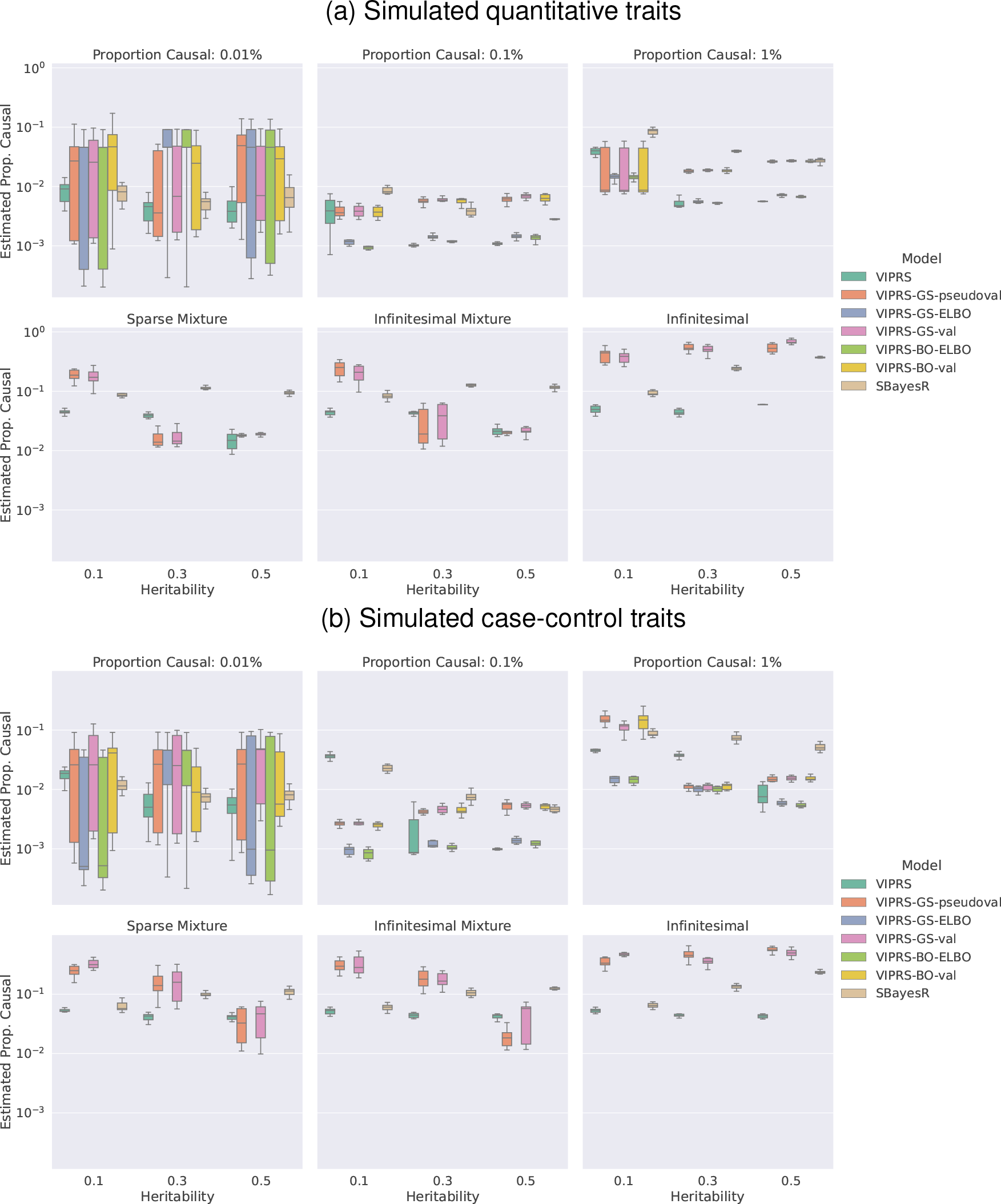
Estimates of polygenicity or proportion of causal variants by summary statistics-based PRS methods for simulated **(a)** quantitative and **(b)** case-control traits. The simulated phenotypes encompass a total of 6 genetic architectures and 3 values for SNP heritability (see **Methods**). Each panel shows results for phenotypes simulated with the pre-specified architecture and each column within a panel shows performance metrics for phenotypes simulated with a pre-specified SNP heritability. The performance metrics are **(a)** incremental prediction *R*^2^ for quantitative traits and **(b)** area under the Precision Recall curve (AUPRC) for binary traits. The boxplot for each method and simulation configuration shows the quartiles of predictive scores for the 10 simulated phenotypes. The methods tested are `VIPRS` (Variational EM), `VIPRS-GS-pseudoval` (grid search with pseudovalidation), `VIPRS-GS-val` (grid search with validation), `VIPRS-BO-ELBO` (Bayesian optimization with the ELBO as criterion), `VIPRS-BO-val` (Bayesian optimization with validation), and `SBayesR`.

**Figure S11:**
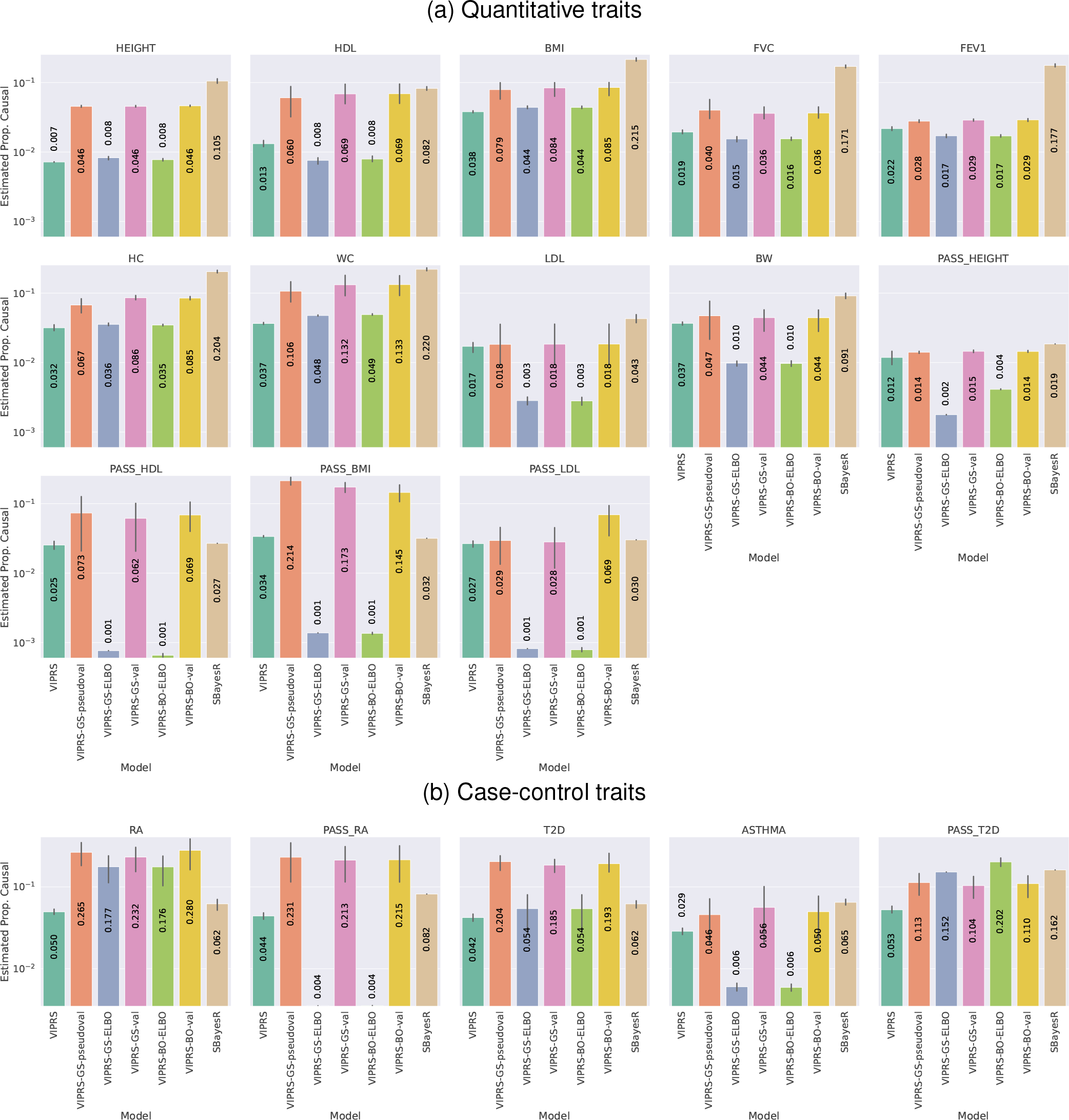
Estimates of polygenicity or proportion of causal variants by summary statistics-based PRS methods for real **(a)** quantitative and **(b)** case-control traits in the UK Biobank. The bars show the mean estimate of polygenicity across the 5 folds and the black lines show the corresponding standard errors. The phenotypes analyzed are described in detail in Table 1. The methods tested are `VIPRS` (Variational EM), `VIPRS-GS-pseudoval` (grid search with pseudovalidation), `VIPRS-GS-val` (grid search with validation), `VIPRS-BO-ELBO` (Bayesian optimization with the ELBO as criterion), `VIPRS-BO-val` (Bayesian optimization with validation), and `SBayesR`.

#### S2.6 Correlation between polygenic score estimates of different PRS methods

**Figure S12:**
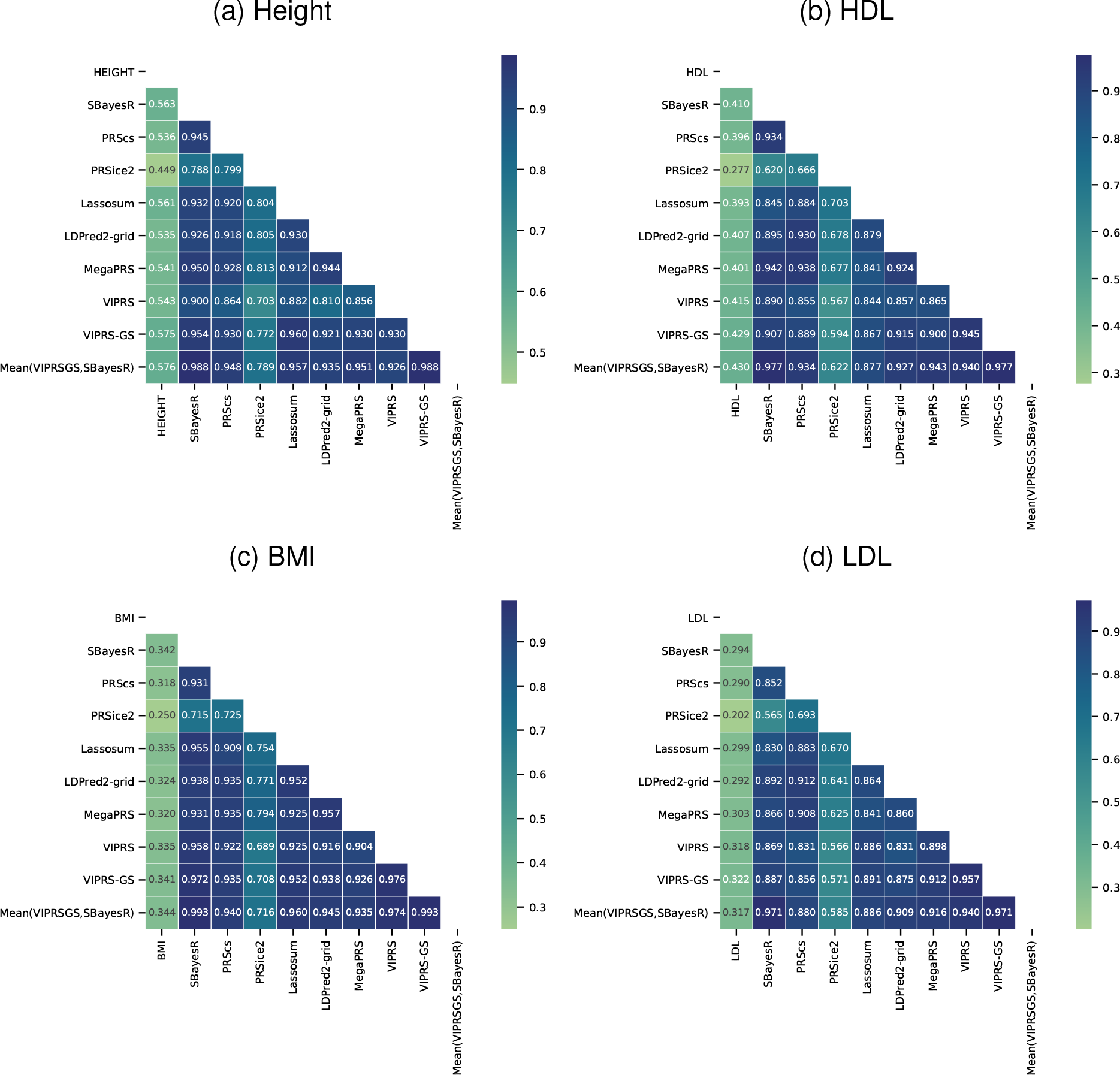
Pearson correlation coefficient between PRS estimates for the UK Biobank individuals derived from different summary statistics-based PRS methods for **(a)** standing height (HEIGHT) **(b)** High-Density Lipoprotein (HDL) and **(c)** body mass index (BMI) and **(d)** Low-Density Lipoprotein (LDL). Each panel shows a heatmap for a different phenotype and each cell shows the Pearson correlation coefficient between the polygenic score estimates of a pair of PRS methods. In addition to the phenotype itself, we show the PRS correlations between `SBayesR`, `PRScs`, `PRSice2`, `Lassosum`, LDPred2-grid, `MegaPRS`, `VIPRS`, `VIPRS-GS`, and Mean(`VIPRSGS`,`SBayesR`) (the averaged PRS estimate between `SBayesR` and `VIPRS-GS`).

#### S2.7 Relationship between the ELBO and validation R^2^ in analyses with real quantitative traits

**Figure S13:**
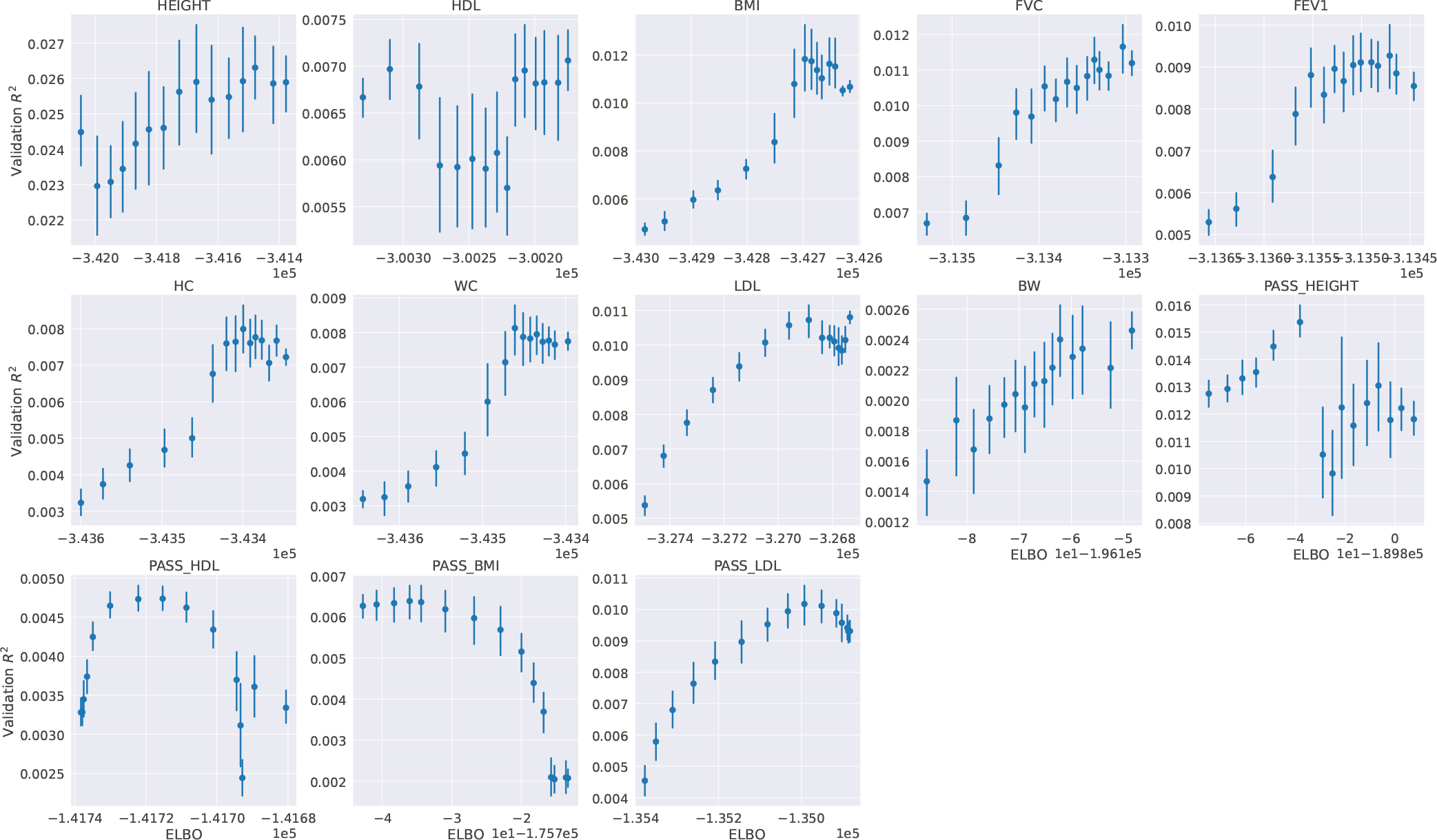
The correspondence between the Evidence Lower BOund (ELBO) of the `VIPRS-GS` models at convergence and the prediction *R*^2^ on the held-out validation set in the UK Biobank. When we train the `VIPRS-GS` version of the model, we are in effect training 30 separate `VIPRS` models with different *π* values. At convergence, we store the maximum value for the ELBO as well as the prediction *R*^2^ on the validation set. Here, we compare the ELBO value at convergence to the corresponding validation *R*^2^. The phenotypes analyzed are described in detail in Table 1. For readability, for each phenotype and/or GWAS data source (panel), the values for the ELBO on the x-axis were grouped into 15 bins. The dots show the mean validation *R*^2^ within that bin and across the 5 training folds and the vertical lines show the corresponding standard errors. Since we perform model fit on each chromosome separately, this figure shows the metrics from the model training with chromosome 2.

## Footnotes

1 This behavior has been partially documented in the context of the `SBayesR` model. See ["Achieving SBayesR convergence for Locke et al. 2015 - BMI GWAS and Wood et al. 2014 - Height GWAS."](https://www.dropbox.com/sh/9aougeoxw4ygo8k/AAD6PT3a3ggv1-KYHytbeUNha?dl=0&preview=SBayesR_locke_wood_convergence.docx).

## Notes

### Competing Interest Statement

The authors have declared no competing interest.

### Summary of Updates

New results

